# OpenPBTA: An Open Pediatric Brain Tumor Atlas

**DOI:** 10.1101/2022.09.13.507832

**Authors:** Joshua A. Shapiro, Krutika S. Gaonkar, Candace L. Savonen, Stephanie J. Spielman, Chante J. Bethell, Run Jin, Komal S. Rathi, Yuankun Zhu, Laura E. Egolf, Bailey K. Farrow, Daniel P. Miller, Yang Yang, Tejaswi Koganti, Nighat Noureen, Mateusz P. Koptyra, Nhat Duong, Adam A. Kraya, Cassie Kline, Yiran Guo, Phillip B. Storm, Javad Nazarian, Stephen C. Mack, Pichai Raman, Siyuan Zheng, Peter J. Madsen, Nicholas Van Kuren, Shannon Robins, Xiaoyan Huang, Angela J. Waanders, Jung Kim, Derek Hanson, Carl J. Koschmann, Sharon J. Diskin, Rishi R. Lulla, Miguel A. Brown, Jessica Wong, Jennifer L. Mason, Laura Scolaro, Meen Chul Kim, Hongbo M. Xie, Brian R. Rood, Kristina A. Cole, Ashley S. Margol, Zalman Vaksman, Rebecca S. Kaufman, Allison P. Heath, Brian M. Ennis, Mariarita Santi, Anna R. Poetsch, Angela N. Viaene, Douglas R. Stewart, Steven M. Foltz, Payal Jain, Bo Zhang, Shrivats Kannan, Michael Prados, Jena V. Lilly, Sabine Mueller, Adam C. Resnick, Casey S. Greene, Jo Lynne Rokita, Jaclyn N. Taroni, Children’s Brain Tumor Network, Pacific Pediatric Neuro-Oncology Consortium

**Affiliations:** Childhood Cancer Data Lab, Alex’s Lemonade Stand Foundation, Bala Cynwyd, PA, USA · Funded by Alex’s Lemonade Stand Foundation Childhood Cancer Data Lab (CCDL); Center for Data-Driven Discovery in Biomedicine, Children’s Hospital of Philadelphia; Division of Neurosurgery, Children’s Hospital of Philadelphia; Department of Bioinformatics and Health Informatics, Children’s Hospital of Philadelphia; Cell and Molecular Biology Graduate Group, Perelman School of Medicine at the University of Pennsylvania; Division of Oncology, Children’s Hospital of Philadelphia; Ben May Department for Cancer Research, University of Chicago, Chicago IL, USA; Greehey Children’s Cancer Research Institute, UT Health San Antonio; Division of Oncology, Children’s Hospital of Philadelphia; Center for Data-Driven Discovery in Biomedicine, Children’s Hospital of Philadelphia; Division of Neurosurgery, Children’s Hospital of Philadelphia · Funded by Alex’s Lemonade Stand Foundation (Catalyst); Children’s Hospital of Philadelphia Division of Neurosurgery; Children’s National Research Institute, Washington,D.C.; George Washington University School of Medicine and Health Sciences, Washington, D.C.; Department of Developmental Neurobiology, St. Jude Children’s Research Hospital; Division of Neurosurgery, Children’s Hospital of Philadelphia; Center for Data-Driven Discovery in Biomedicine, Children’s Hospital of Philadelphia; Division of Hematology, Oncology, Neuro-Oncology, and Stem Cell Transplant, Ann & Robert H Lurie Children’s Hospital of Chicago; Department of Pediatrics, Northwestern University Feinberg School of Medicine; Clinical Genetics Branch, Division of Cancer Epidemiology and Genetics, National Cancer Institute; Department of Pediatrics, University of Michigan Health, Ann Arbor, MI; Pediatric Hematology Oncology,Mott Children’s Hospital, Ann Arbor, MI; Division of Hematology/Oncology, Hasbro Children’s Hospital; Department of Pediatrics, The Warren Alpert School of Brown University, Providence, Rhode Island; Department of Pediatrics, University of Pennsylvania, Philadelphia, PA; Abramson Family Cancer Research Institute, Perelman School of Medicine at the University of Pennsylvania, Philadelphia, PA; Division of Hematology and Oncology, Children’s Hospital Los Angeles; Department of Pediatrics, Keck School of Medicine of University of Southern California; Center for Data-Driven Discovery in Biomedicine, Children’s Hospital of Philadelphia; Division of Neurosurgery, Children’s Hospital of Philadelphia · Funded by NIH U2C HL138346-03; NCI/NIH Contract No. 75N91019D00024, Task Order No. 75N91020F00003; Australian Government,Department of Education; Department of Pathology and Laboratory Medicine, Children’s Hospital of Philadelphia; Department of Pathology and Laboratory Medicine, University of Pennsylvania Perelman School of Medicine; Biotechnology Center, Technical University Dresden, Germany; National Center for Tumor Diseases, Dresden, Germany · Funded by The St. Anna Kinderkrebsforschung,Austria; The Mildred Scheel Early Career Center Dresden P2, funded by the German Cancer Aid; Department of Systems Pharmacology and Translational Therapeutics, University of Pennsylvania; Childhood Cancer Data Lab, Alex’s Lemonade Stand Foundation, Bala Cynwyd, PA, USA · Funded by Alex’s Lemonade Stand Foundation GR-000002471; National Institutes of Health K12GM081259; University of California, San Francisco, San Francisco, CA, USA; Center for Data-Driven Discovery in Biomedicine, Children’s Hospital of Philadelphia; Division of Neurosurgery, Children’s Hospital of Philadelphia · Funded by Alex’s Lemonade Stand Foundation (Catalyst); Children’s Brain Tumor Network; NIH 3P30 CA016520-44S5, U2C HL138346-03, U24 CA220457-03; NCI/NIH Contract No. 75N91019D00024,Task Order No. 75N91020F00003; Children’s Hospital of Philadelphia Division of Neurosurgery; Department of Systems Pharmacology and Translational Therapeutics, Perelman School of Medicine, University of Pennsylvania, Philadelphia, PA, USA; Childhood Cancer Data Lab, Alex’s Lemonade Stand Foundation, Bala Cynwyd, PA, USA; Center for Health AI, University of Colorado School of Medicine, Aurora, CO, USA; Center for Data-Driven Discovery in Biomedicine, Children’s Hospital of Philadelphia; Division of Neurosurgery, Children’s Hospital of Philadelphia; Department of Bioinformatics and Health Informatics, Children’s Hospital of Philadelphia · Funded by Alex’s Lemonade Stand Foundation (Young Investigator, Catalyst); NCI/NIH Contract No. 75N91019D00024, Task Order No. 75N91020F00003; Hackensack Meridian 93 School of Medicine; Hackensack University Medical Center

**Keywords:** pediatric cancer, brain tumors, somatic variation, open science, reproducibility, classification, tumor atlas

## Abstract

**Summary:** Pediatric brain and spinal cancer are the leading disease-related cause of death in children, thus we urgently need curative therapeutic strategies for these tumors. To accelerate such discoveries, the Children’s Brain Tumor Network and Pacific Pediatric Neuro-Oncology Consortium created a systematic process for tumor biobanking, model generation, and sequencing with immediate access to harmonized data. We leverage these data to create OpenPBTA, an open collaborative project which establishes over 40 scalable analysis modules to genomically characterize 1,074 pediatric brain tumors. Transcriptomic classification reveals that *TP53* loss is a significant marker for poor overall survival in ependymomas and H3 K28-altered diffuse midline gliomas and further identifies universal *TP53* dysregulation in mismatch repair-deficient hypermutant high-grade gliomas. OpenPBTA is a foundational analysis platform actively being applied to other pediatric cancers and inform molecular tumor board decision-making, making it an invaluable resource to the pediatric oncology community.

**In Brief:** The OpenPBTA is a global, collaborative open-science initiative which brought together researchers and clinicians to genomically characterize 1,074 pediatric brain tumors and 22 patient-derived cell lines. Shapiro, et. al create over 40 open-source, scalable modules to perform cancer genomics analyses and provide a richly-annotated somatic dataset across 58 brain tumor histologies. The OpenPBTA framework can be used as a model for large-scale data integration to inform basic research, therapeutic target identification, and clinical translation.

**Highlights:** OpenPBTA collaborative analyses establish resource for 1,074 pediatric brain tumors NGS-based WHO-aligned integrated diagnoses generated for 641 of 1,074 tumors RNA-Seq analysis infers medulloblastoma subtypes, TP53 status, and telomerase activity OpenPBTA will accelerate therapeutic translation of genomic insights

## Introduction

Pediatric brain and spinal cord tumors are collectively the second most common malignancy in children after leukemia, and they represent the leading disease-related cause of death in children^1^. Five-year survival rates vary widely across different histologic and molecular classifications of brain tumors. For example, most high-grade gliomas carry a universally fatal prognosis, while children with pilocytic astrocytoma have an estimated 10-year survival rate of 92%^2^. Moreover, estimates from 2009 suggest that children and adolescents aged 0-19 with brain tumors in the United States have lost an average of 47,631 years of potential life^3^.

The low survival rates for some pediatric tumors are clearly multifactorial, explained partly by our lack of comprehensive understanding of the ever-evolving array of brain tumor molecular subtypes, difficulty drugging these tumors, and the shortage of drugs specifically labeled for pediatric malignancies. Historically, some of the most fatal, inoperable brain tumors, such as diffuse intrinsic pontine gliomas (DIPGs), were not routinely biopsied due to perceived risks of biopsy and the paucity of therapeutic options that would require tissue. Limited access to tissue to develop patient-derived cell lines and mouse models has been a barrier to research. Furthermore, the incidence of any single brain tumor molecular subtype is relatively low due to the rarity of pediatric tumors in general.

To address these long-standing barriers, multiple national and international consortia have come together to uniformly collect clinically-annotated surgical biosamples and associated germline materials as part of both observational and interventional clinical trials. Such accessible, centralized resources enable collaborative sharing of specimens and data across rare cancer subtypes to accelerate breakthroughs and clinical translation. The creation of the Pediatric Brain Tumor Atlas (PBTA) in 2018, led by the Children’s Brain Tumor Network (CBTN, cbtn.org) and the Pacific Pediatric Neuro-Oncology Consortium (PNOC, PNOC.us) is one such effort that builds on nearly 10 years of multi-institutional enrollment, sample collection, and clinical followup across more than 30 institutions. Just as cooperation is required to share specimens and data, rigorous cancer genomic analysis requires collaboration among researchers with distinct expertise, such as computational scientists, bench scientists, clinicians, and pathologists.

Although there has been significant progress in recent years to elucidate the landscape of somatic variation responsible for pediatric brain tumor formation and progression, translation of therapeutic agents to phase II or III clinical trials and subsequent FDA approvals have not kept pace. Within the last 20 years, the FDA has approved only five drugs for the treatment of pediatric brain tumors: mTOR inhibitor everolimus, for subependymal giant cell astrocytoma; anti-PD-1 immunotherapy pembrolizumab, for microsatellite instability–high or mismatch repair– deficient tumors; NTRK inhibitors larotrectinib and entrectinib, for tumors with an NTRK 1/2/3 gene fusions; and MEK1/2 inhibitor selumetinib, for neurofibromatosis type 1 (NF1) and symptomatic, inoperable plexiform neurofibromas^4^.

This is, in part, due to pharmaceutical company priorities and concerns regarding toxicity, making it challenging for researchers to obtain to new therapeutic agents for pediatric clinical trials. Critically, as of August 18, 2020, an amendment to the Pediatric Research Equity Act called the “Research to Accelerate Cures and Equity (RACE) for Children Act” mandates that all new adult oncology drugs also be tested in children when the molecular targets are relevant to a particular childhood cancer. The regulatory change introduced by the RACE Act, coupled with the identification of putative molecular targets in pediatric cancers through genomic characterization, is poised to accelerate identification of novel and effective therapeutic for pediatric diseases that have otherwise been overlooked.

To leverage diverse scientific and analytical expertise to analyze the PBTA data, we created an open science model and incorporated features such as analytical code review^5, 6^ and continuous integration to test data and code^6, 7^ to improve reproducibility throughout the life cycle of our project, termed OpenPBTA.

We anticipated that a model of open collaboration would enhance the value of our effort to the pediatric brain tumor research community and provide a framework for continuous, accelerated translation of pediatric brain tumor datasets. Openly sharing data and code in real time allows others to build upon the work more rapidly, and publications that include data and code sharing are poised for greater impact^8, 9^. Here, we present a comprehensive, collaborative, open genomic analysis of 1,074 tumors and 22 cell lines, comprised of 58 distinct brain tumor histologies from 943 patients. The data and containerized infrastructure of OpenPBTA have been instrumental for discovery and translational research studies^10–12^, are actively integrated into PNOC molecular tumor board decision-making, and are a foundational layer for the NCI’s Childhood Cancer Data Initiative’s (CCDI) pediatric Molecular Targets Platform (https://moleculartargets.ccdi.cancer.gov/) recently built in support of the RACE Act^13^. We anticipate OpenPBTA will be an invaluable resource to the pediatric oncology community.

## Results

### Crowd-sourced Somatic Analyses to Create an Open Pediatric Brain Tumor Atlas

We previously performed whole genome sequencing (WGS), whole exome sequencing (WXS), and RNA sequencing (RNA-Seq) on matched tumor and normal tissues as well as selected cell lines^14^ from 943 patients from the Pediatric Brain Tumor Atlas (PBTA), consisting of samples from the <u>Children’s Brain Tumor Network (CBTN)</u> and the PNOC003 DIPG clinical trial^12, 15^ of the <u>Pacific Pediatric Neuro-Oncology Consortium (PNOC)</u> (Figure 1A). We then harnessed the benchmarking efforts of the <u>Gabriella Miller Kids First Data Resource Center</u> to develop robust and reproducible data analysis workflows within the CAVATICA platform to perform primary somatic analyses including calling of single nucleotide variants (SNVs), copy number variants (CNVs), structural variants (SVs), and gene fusions, often implementing multiple complementary methods (Figure S1) and **STAR Methods**).

To facilitate analysis and visualization of this large, diverse cohort, we further categorized tumor broad histologies (i.e., broad 2016 WHO classifications) into smaller groupings we denote “cancer groups.” A summarized view of the number of biospecimens per phase of therapy across different broad histologies and cancer groups is shown in (Figure 1B). We maintained a data release folder on Amazon S3, downloadable directly from S3 or through the open-access <u>CAVATICA project</u>, with merged files for each analysis (See **Data and code availability** section). As new analytical results (e.g., tumor mutation burden calculations) that we expected to be used across multiple analyses were produced, or issues with the data were identified, new data releases were made available in a versioned manner.

A key innovative feature of this project has been its open contribution model used for both analyses (i.e., analytical code) and scientific manuscript writing. We created a public Github analysis repository (https://github.com/AlexsLemonade/OpenPBTA-analysis) to hold all code associated with analyses downstream of the Kids First Data Resource Center workflows and a GitHub manuscript repository (https://github.com/AlexsLemonade/OpenPBTA-manuscript) with Manubot^16^ integration to enable real-time manuscript creation using Markdown within GitHub. Importantly, all analyses and manuscript writing were conducted openly throughout the research project, allowing any researcher in the world the opportunity to contribute.

The process for analysis and manuscript contributions is outlined in Figure 1C. First, a potential contributor would propose an analysis by filing an issue in the GitHub analysis repository. Next, organizers for the project, or other contributors with expertise, had the opportunity to provide feedback about the proposed analysis (Figure 1C). The contributor then made a copy (fork) of the analysis repository and added their proposed analysis code and results to their fork. The contributor would formally request to include their analytical code and results to the main OpenPBTA analysis repository by filing a pull request on GitHub. All pull requests to the analysis repository underwent peer review by organizers and/or other contributors to ensure scientific accuracy, maintainability, and readability of code and documentation (Figure 1C-D).

The collaborative nature of the project required additional steps beyond peer review of analytical code to ensure consistent results for all collaborators and over time (Figure 1D). We leveraged Docker®^17^ and the Rocker project^18^ to maintain a consistent software development environment, creating a monolithic image that contained all dependencies necessary for analyses. To ensure that new code would execute in the development environment, we used the continuous integration (CI) service CircleCI® to run analytical code on a small subset of data for testing before formal code review, allowing us to detect code bugs or sensitivity to changes in the underlying data.

We followed a similar process in our Manubot-powered^16^ manuscript repository for additions to the manuscript (Figure 1C). Contributors forked the manuscript repository, added proposed content to their branch, and filed pull requests to the main manuscript repository with their changes. Similarly, pull requests underwent a peer review process for clarity and correctness, agreement with interpretation, and spell checking via Manubot.

**Figure 1.**
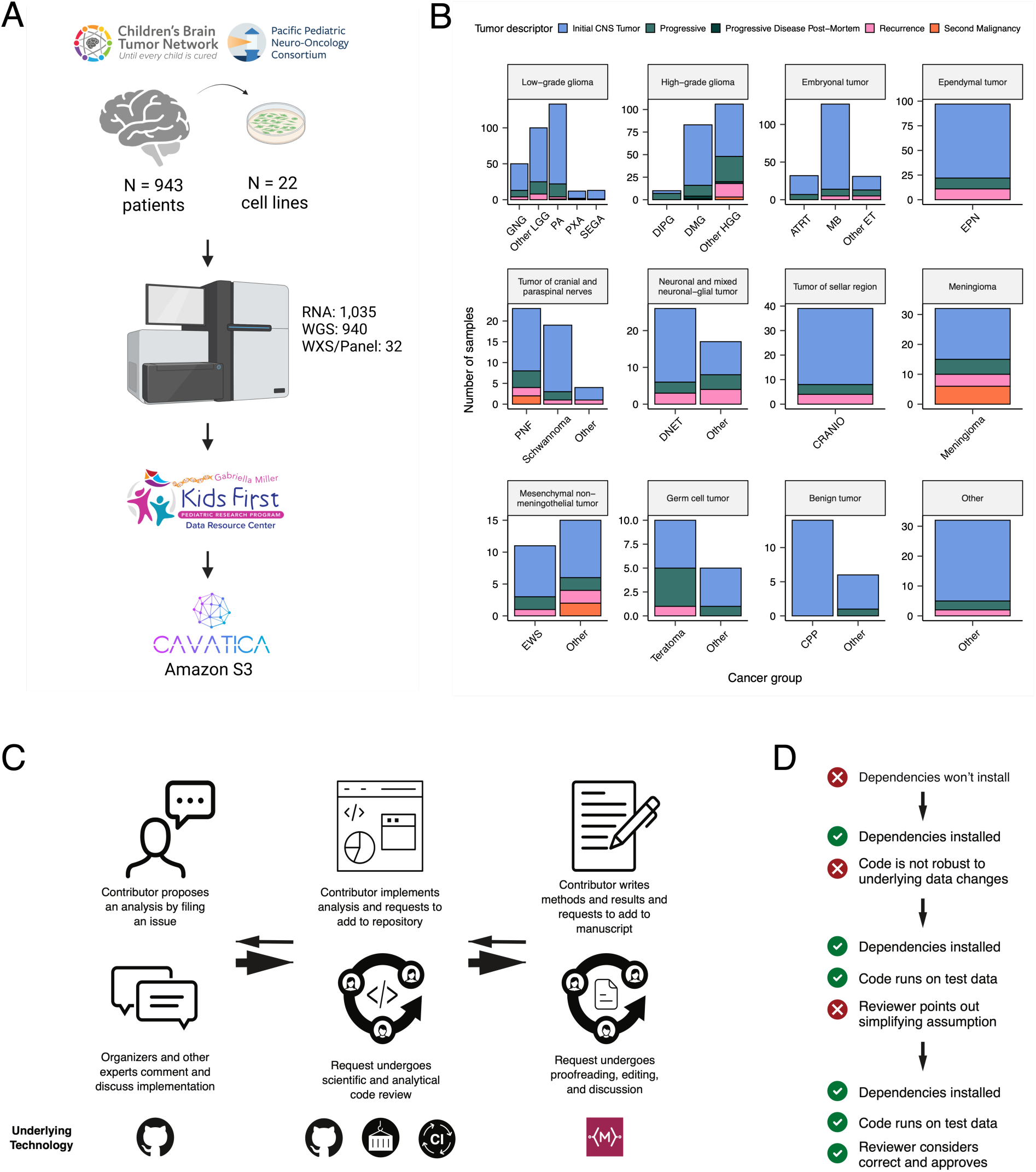
Overview of the OpenPBTA Project. A, The Children’s Brain Tumor Network and the Pacific Pediatric Neuro-Oncology Consortium collected tumor samples from 943 patients. To date, 22 cell lines were created from tumor tissue, and over 2000 specimens were sequenced (N = 1035 RNA-Seq, N = 940 WGS, and N = 32 WXS or targeted panel). Data was harmonized by the Kids First Data Resource Center using an Amazon S3 framework within CAVATICA. B, Stacked bar plot summary of the number of biospecimens per phase of therapy per broad histology (Abbreviations: GNG = ganglioglioma, Other LGG = other low-grade glioma, PA = pilocytic astrocytoma, PXA = pleomorphic xanthoastrocytoma, SEGA = subependymal giant cell astrocytoma, DIPG = diffuse intrinsic pontine glioma, DMG = diffuse midline glioma, Other HGG = other high-grade glioma, ATRT = atypical teratoid rhabdoid tumor, MB = medulloblastoma, Other ET = other embryonal tumor, EPN = ependymoma, PNF = plexiform neurofibroma, DNET = dysembryoplastic neuroepithelial tumor, CRANIO = craniopharyngioma, EWS = Ewing sarcoma, CPP = choroid plexus papilloma). Only samples with available descriptors were included. C, Overview of the open analysis and manuscript contribution model. In the analysis GitHub repository, a contributor would propose an analysis that other participants can comment on. Contributors would then implement the analysis and file a request to add their changes to the analysis repository (“pull request”). Pull requests underwent review for scientific rigor and correctness of implementation. Pull requests were additionally checked to ensure that all software dependencies were included and the code was not sensitive to underlying data changes using container and continuous integration technologies. Finally, a contributor would file a pull request documenting their methods and results to the Manubot-powered manuscript repository. Pull requests in the manuscript repository were also subject to review. D, A potential path for an analytical pull request. Arrows indicate revisions to a pull request. Prior to review, a pull request was tested for dependency installation and whether or not the code would execute. Pull requests also required approval by organizers and/or other contributors, who checked for scientific correctness. Panel A created with BioRender.com.

### Molecular Subtyping of OpenPBTA CNS Tumors

Over the past two decades, experts in neuro-oncology have worked with the World Health Organization (WHO) to iteratively redefine the classifications of central nervous system (CNS) tumors^19, 20^. More recently, in 2016 and 2021^21, 22^, molecular subtypes have been integrated into these classifications. In 2011, the Children’s Brain Tumor Tissue Consortium, now known as the Children’s Brain Tumor Network (CBTN), opened its protocol for brain tumor and matched normal sample collection. Since the CBTN opened its collection protocol in 2011, before molecular data were integrated into classifications, the majority of the samples within the OpenPBTA lacked molecular subtype annotations at the time of tissue collection. Moreover, the OpenPBTA data does not yet feature methylation arrays which are increasingly used to inform molecular subtyping. Therefore, we jointly considered key genomic features of tumor entities described by the WHO in 2016, low-grade glioma (LGG) subtypes described by Ryall and colleagues^23^, as well as clinician and pathologist review, to generate research-grade integrated diagnoses for 60% (641/1074) of tumor samples with high confidence (**Table S1**).

Importantly, this collaborative molecular subtyping process allowed us to identify potential data entry errors (e.g., an ETMR incorrectly entered as a medulloblastoma) and histologically mis-identified specimens (e.g., Ewing sarcoma sample labeled as a craniopharyngioma), update diagnoses using current WHO terms (e.g., tumors formerly ascribed primitive neuro-ectodermal tumor [PNET] diagnoses), and discover rarer tumor entities within the OpenPBTA (e.g., H3-mutant ependymoma, meningioma with *YAP1::FAM118B* fusion). Table 1 lists the subtypes we defined within OpenPBTA, comprising low-grade gliomas (N = 290), high-grade gliomas (N = 141), embryonal tumors (N = 126), ependymomas (N = 30), tumors of sellar region (N = 27), mesenchymal non-meningothelial tumors (N = 11), glialneuronal tumors (N = 10), and chordomas (N = 6). For detailed methods, see **STAR Methods** and Figure S1.

**Table 1.** Molecular subtypes generated through the OpenPBTA project. Listed are broad tumor histologies, molecular subtypes generated, and number of specimens subtyped within the OpenPBTA project.

| Broad histology group | OpenPBTA molecular subtype | n |
| --- | --- | --- |
| Chordoma | CHDM, conventional | 2 |
| Chordoma | CHDM, poorly differentiated | 4 |
| Embryonal tumor | CNS Embryonal, NOS | 13 |
| Embryonal tumor | CNS HGNET-MN1 | 1 |
| Embryonal tumor | CNS NB-FOXR2 | 3 |
| Embryonal tumor | ETMR, C19MC-altered | 5 |
| Embryonal tumor | ETMR, NOS | 1 |
| Embryonal tumor | MB, Group3 | 14 |
| Embryonal tumor | MB, Group4 | 49 |
| Embryonal tumor | MB, SHH | 30 |
| Embryonal tumor | MB, WNT | 10 |
| Ependymal tumor | EPN, H3 K28 | 1 |
| Ependymal tumor | EPN, ST RELA | 28 |
| Ependymal tumor | EPN, ST YAP1 | 1 |
| High-grade glioma | DMG, H3 K28 | 24 |
| High-grade glioma | DMG, H3 K28, TP53 activated | 13 |
| High-grade glioma | DMG, H3 K28, TP53 loss | 40 |
| High-grade glioma | HGG, H3 G35 | 3 |
| High-grade glioma | HGG, H3 G35, TP53 loss | 1 |
| High-grade glioma | HGG, H3 wildtype | 31 |
| High-grade glioma | HGG, H3 wildtype, TP53 activated | 5 |
| High-grade glioma | HGG, H3 wildtype, TP53 loss | 21 |
| High-grade glioma | HGG, IDH, TP53 activated | 2 |
| High-grade glioma | HGG, IDH, TP53 loss | 1 |
| Low-grade glioma | GNG, BRAF V600E | 13 |
| Low-grade glioma | GNG, BRAF V600E, CDKN2A/B | 1 |
| Low-grade glioma | GNG, FGFR | 1 |
| Low-grade glioma | GNG, H3 | 1 |
| Low-grade glioma | GNG, IDH | 2 |
| Low-grade glioma | GNG, KIAA1549-BRAF | 5 |
| Low-grade glioma | GNG, MYB/MYBL1 | 1 |

| **Broad histology group** | **OpenPBTA molecular subtype** | **n** |
| --- | --- | --- |
| Low-grade glioma | GNG, NF1-germline | 1 |
| Low-grade glioma | GNG, NF1-somatic, BRAF V600E | 1 |
| Low-grade glioma | GNG, other MAPK | 4 |
| Low-grade glioma | GNG, other MAPK, IDH | 1 |
| Low-grade glioma | GNG, RTK | 3 |
| Low-grade glioma | GNG, wildtype | 14 |
| Low-grade glioma | LGG, BRAF V600E | 27 |
| Low-grade glioma | LGG, BRAF V600E, CDKN2A/B | 5 |
| Low-grade glioma | LGG, FGFR | 8 |
| Low-grade glioma | LGG, IDH | 3 |
| Low-grade glioma | LGG, KIAA1549-BRAF | 113 |
| Low-grade glioma | LGG, KIAA1549-BRAF, NF1-germline | 1 |
| Low-grade glioma | LGG, KIAA1549-BRAF, other MAPK | 1 |
| Low-grade glioma | LGG, MYB/MYBL1 | 2 |
| Low-grade glioma | LGG, NF1-germline | 6 |
| Low-grade glioma | LGG, NF1-germline, CDKN2A/B | 1 |
| Low-grade glioma | LGG, NF1-germline, FGFR | 2 |
| Low-grade glioma | LGG, NF1-somatic | 2 |
| Low-grade glioma | LGG, NF1-somatic, FGFR | 1 |
| Low-grade glioma | LGG, NF1-somatic, NF1-germline, CDKN2A/B | 1 |
| Low-grade glioma | LGG, other MAPK | 12 |
| Low-grade glioma | LGG, RTK | 10 |
| Low-grade glioma | LGG, RTK, CDKN2A/B | 1 |
| Low-grade glioma | LGG, wildtype | 34 |
| Low-grade glioma | SEGA, RTK | 1 |
| Low-grade glioma | SEGA, wildtype | 11 |
| Mesenchymal non-meningothelial tumor | EWS | 11 |
| Neuronal and mixed neuronal-glial tumor | CNC | 2 |
| Neuronal and mixed neuronal-glial tumor | EVN | 1 |
| Neuronal and mixed neuronal-glial tumor | GNT, BRAF V600E | 1 |
| Neuronal and mixed neuronal-glial tumor | GNT, KIAA1549-BRAF | 2 |
| Neuronal and mixed neuronal-glial tumor | GNT, other MAPK | 1 |
| Neuronal and mixed neuronal-glial tumor | GNT, other MAPK, FGFR | 1 |
| Neuronal and mixed neuronal-glial tumor | GNT, RTK | 2 |
| Tumor of sellar region | CRANIO, ADAM | 27 |
|  | Total | 641 |

### Somatic Mutational Landscape of Pediatric Brain Tumors

We performed a comprehensive genomic analysis of somatic SNVs, CNVs, SVs, and fusions across 1,074 tumors (N = 1,019 RNA-Seq, N = 918 WGS, N = 32 WXS/Panel) and 22 cell lines (N = 16 RNA-Seq, N = 22 WGS), from 943 patients, 833 with paired normal specimens (N = 801 WGS, N = 32 WXS/Panel). Following SNV consensus calling (Figure S1 and Figure S2A-G), we observed as expected lower tumor mutation burden (TMB) Figure S2H in pediatric tumors compared to adult brain tumors from The Cancer Genome Atlas (TCGA), Figure S2I, with hypermutant (> 10 Mut/Mb) and ultra-hypermutant (> 100 Mut/Mb) tumors^24^ only found within HGGs. Figure 2 and Figure S3A depict oncoprints of histology-specific driver genes across PBTA histologies.

### Low-grade gliomas

As expected, the majority (62%, 140/227) of LGGs harbored a somatic alteration in *BRAF*, with canonical *BRAF::KIAA1549* fusions as the major oncogenic driver^25^ (Figure 2A). We observed additional mutations in *FGFR1* (2%), *PIK3CA* (2%), *KRAS* (2%), *TP53* (1%), and *ATRX* (1%) and fusions in *NTRK2* (2%), *RAF1* (2%), *MYB* (1%), *QKI* (1%), *ROS1* (1%), and *FGFR2* (1%), concordant with previous studies reporting the near universal upregulation of the RAS/MAPK pathway in these tumors resulting from activating mutations and/or oncogenic fusions^23, 25^. Indeed, we observed significant upregulation (ANOVA Bonferroni-corrected p < 0.01) of the KRAS signaling pathway in LGGs (Figure 5B).

### Embryonal tumors

The majority (N = 95) of embryonal tumor samples were medulloblastomas that spanned the spectrum of molecular subtypes (WNT, SHH, Group3, and Group 4; see **Molecular Subtyping of CNS Tumors**), as identified by subtype-specific canonical mutations (Figure 2B). We detected canonical *SMARCB1/SMARCA4* deletions or inactivating mutations in atypical teratoid rhabdoid tumors (ATRTs) and C19MC amplification in the embryonal tumors with multilayer rosettes (ETMRs, displayed as other embryonal tumors)^26–29^.

### High-grade gliomas

Across HGGs, we found that *TP53* (57%, 35/61) and *H3F3A* (52%, 32/61) were both most mutated and co-occurring genes (Figure 2A and C), followed by frequent mutations in *ATRX* (30%, 18/61). We observed recurrent amplifications and fusions in *EGFR*, *MET*, *PDGFRA*, and *KIT*, highlighting that these tumors utilize multiple oncogenic mechanisms to activate tyrosine kinases, as has been previously reported^15, 30, 31^. Gene set enrichment analysis showed upregulation (ANOVA Bonferroni-corrected p < 0.01) of DNA repair, G2M checkpoint, and MYC pathways as well as downregulation of the TP53 pathway (Figure 5B). The two tumors with ultra-high TMB (> 100 Mutations/Mb) were from patients with known mismatch repair deficiency syndrome^14^.

### Other CNS tumors

We observed that 25% (15/60) of ependymoma tumors were *C11orf95::RELA* (now, *ZFTA::RELA*) fusion-positive ependymomas and that 68% (21/31) of craniopharyngiomas were driven by mutations in *CTNNB1* (Figure 2D). Multiple histologies contained somatic mutations or fusions in *NF2*: 41% (7/17) of meningiomas, 5% (3/60) of ependymomas, and 27% (3/11) schwannomas. Rare fusions in *ERBB4*, *YAP1*, *KRAS*, and *MAML2* were observed in 10% (6/60) of ependymoma tumors. DNETs harbored alterations in MAPK/PI3K pathway genes as previously reported^32^, including *FGFR1* (21%, 4/19), *PDGFRA* (10%, 2/19), and *BRAF* (5%, 1/19). Frequent mutations in additional rare brain tumor histologies are depicted in Figure S3A.

**Figure 2.**
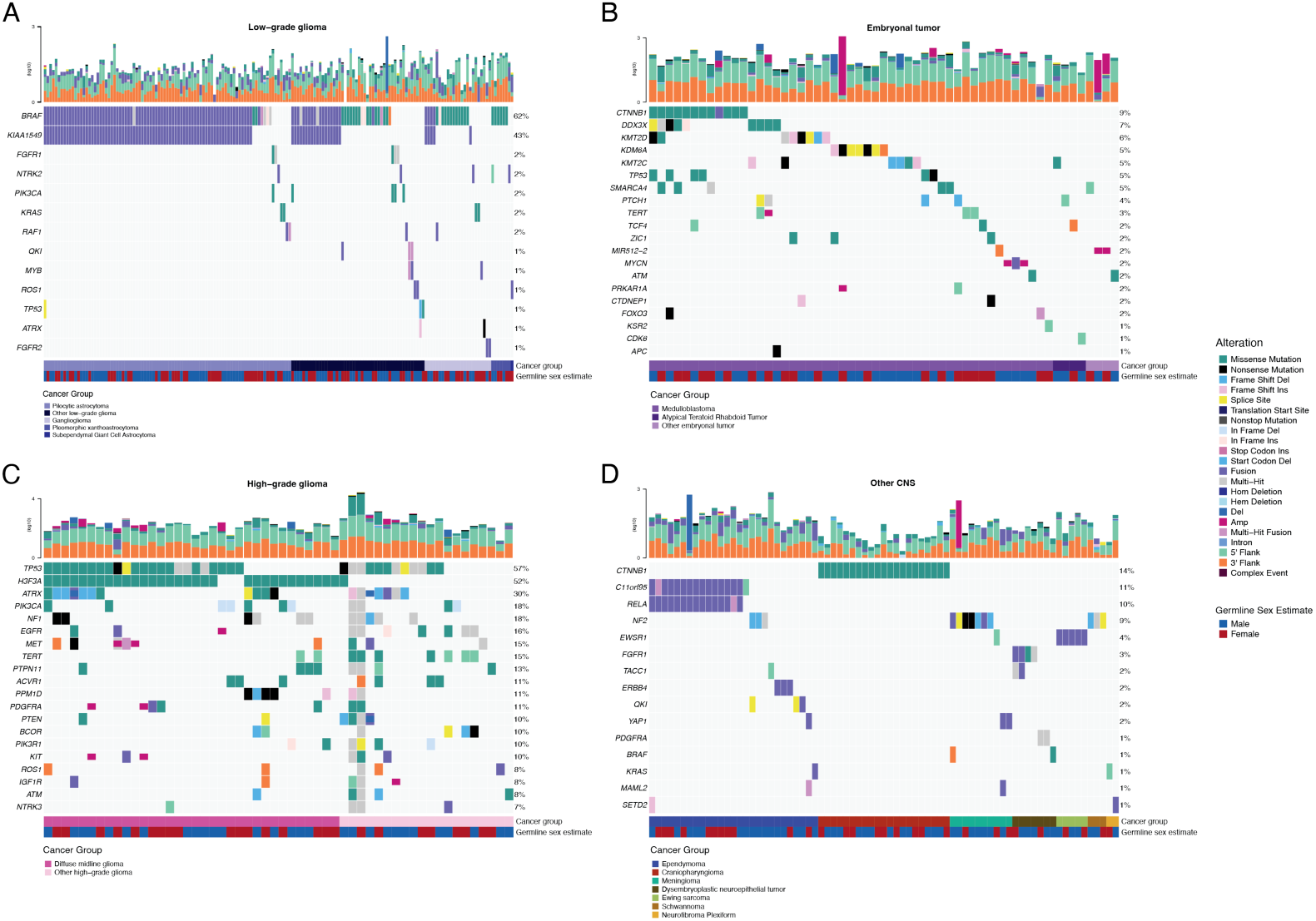
Mutational landscape of PBTA tumors. Shown are frequencies of canonical somatic gene mutations, CNVs, fusions, and TMB (top bar plot) for the top 20 genes mutated across primary tumors within the OpenPBTA dataset. A, Low-grade astrocytic tumors (N = 227): pilocytic astrocytoma (N = 104), other low-grade glioma (N = 69), ganglioglioma (N = 35), pleomorphic xanthoastrocytoma (N = 9), subependymal giant cell astrocytoma (N = 10); B, Embryonal tumors (N = 128): medulloblastomas (N = 95), atypical teratoid rhabdoid tumors (N = 24), other embryonal tumors (N = 9); C, Diffuse astrocytic and oligodendroglial tumors (N = 61): diffuse midline gliomas (N = 34) and other high-grade gliomas (N = 27); D, Other CNS tumors (N = 194): ependymomas (N = 60), craniopharyngiomas (N = 31), meningiomas (N = 17), dysembryoplastic neuroepithelial tumors (N = 19), Ewing sarcomas (N = 7), schwannomas (N = 11), and neurofibroma plexiforms (N = 7). Additional, rare CNS tumors are displayed in Figure S3A. Tumor histology (`Cancer Group`) and patient sex (`Germline sex estimate`) are displayed as annotations at the bottom of each plot. Only samples with mutations in the listed genes are shown. Multiple CNVs are denoted as a complex event.

### Mutational co-occurrence, CNV, and signatures highlight key oncogenic drivers

We analyzed mutational co-occurrence among OpenPBTA tumors, using a single sequencing sample from each individual with available WGS (N = 666). The top 50 mutated genes (see **STAR Methods** for details) in primary tumors are shown in Figure 3 by tumor type (**A**, bar plots), with co-occurrence scores illustrated in the heatmap (**B**). *TP53* was the most frequently mutated gene across OpenPBTA tumors (8.4%, 56/666), significantly co-occurring with *H3F3A* (OR = 32, 95% CI: 15.3 - 66.7, q = 8.46e-17), *ATRX* (OR = 20, 95% CI: 8.4 - 47.7, q = 4.43e-8), *NF1* (OR = 8.62, 95% CI: 3.7 - 20.2, q = 5.45e-5), and *EGFR* (OR = 18.2, 95% CI: 5 - 66.5, q = 1.6e-4). Other canonical cancer driver genes that were frequently mutated included *BRAF*, *H3F3A*, *CTNNB1*, *NF1*, *ATRX*, *FGFR1*, and *PIK3CA*.

At the broad histology level, mutations in *CTNNB1* significantly co-occurred with mutations in *TP53* (OR = 42.9, 95% CI: 7 - 261.4, q = 1.63e-3) and *DDX3X* (OR = 21.1, 95% CI: 4.6 - 96.3, q = 4.46e-3) in embryonal tumors. Mutations in *FGFR1* and *PIK3CA* significantly co-occurred in LGGs (OR = 76.1, 95% CI: 9.85 - 588.1, q = 3.26e-3), consistent with previous findings^33, 34^. Of HGG tumors with mutations in *TP53* or *PPM1D*, 52/54 (96.3%) had mutations in only one of these genes (OR = 0.188, 95% CI: 0.04 - 0.94, p = 0.0413, q = 0.0587). This trend recapitulates previous observations that *TP53* and *PPM1D* mutations tend to be mutually exclusive in HGGs^35^.

We summarized broad CNV and SV and observed that HGGs and DMGs, followed by medulloblastomas, had the most unstable genomes (Figure S3B). By contrast, craniopharyngiomas and schwannomas generally lacked somatic CNV. Together, these CNV patterns largely aligned with our estimates of tumor mutational burden (Figure S2H). The breakpoint density estimated from SV and CNV data was significantly correlated across tumors (p = 1.08e-37) (Figure 3C) and as expected, the number of chromothripsis regions called increased as breakpoint density increased (Figure S3B-C). We identified chromothripsis events in 28% (N = 11/39) of diffuse midline gliomas and in 40% (N = 19/48) of other HGGs (non-midline HGGs) (Figure 3D). We also found evidence of chromothripsis in over 15% of sarcomas, PXAs, metastatic secondary tumors, chordomas, glial-neuronal tumors, germinomas, meningiomas, ependymomas, medulloblastomas, ATRTs, and other embryonal tumors, highlighting the genomic instability and complexity of these pediatric brain tumors.

We next assessed the contributions of eight previously identified adult CNS-specific mutational signatures from the RefSig database^36^ across samples (Figure 3E and Figure S4A). Stage 0 and/or 1 tumors characterized by low TMBs (Figure S2H) such as pilocytic astrocytomas, gangliogliomas, other LGGs, and craniopharyngiomas, were dominated by Signature 1 (Figure S4A), which results from the normal process of spontaneous deamination of 5-methylcytosine. Signature N6 is a CNS-specific signature which we observed nearly universally across samples. Drivers of Signature 18, *TP53*, *APC*, *NOTCH1* (found at https://signal.mutationalsignatures.com/explore/referenceCancerSignature/31/drivers), are also canonical drivers of medulloblastoma, and indeed, we observed Signature 18 as the signature with the highest weight in medulloblastoma tumors. Signatures 3, 8, 18, and MMR2 were prevalent in HGGs, including DMGs. Finally, we found that the Signature 1 weight was higher at diagnosis (pre-treatment) and was almost always lower in tumors at later phases of therapy (progression, recurrence, post-mortem, secondary malignancy; Figure S4B). This trend may have resulted from therapy-induced mutations that produced additional signatures (e.g., temozolomide treatment has been suggested to drive Signature 11^37^), subclonal expansion, and/or acquisition of additional driver mutations during tumor progression, leading to higher overall TMBs and additional signatures.

**Figure 3.**
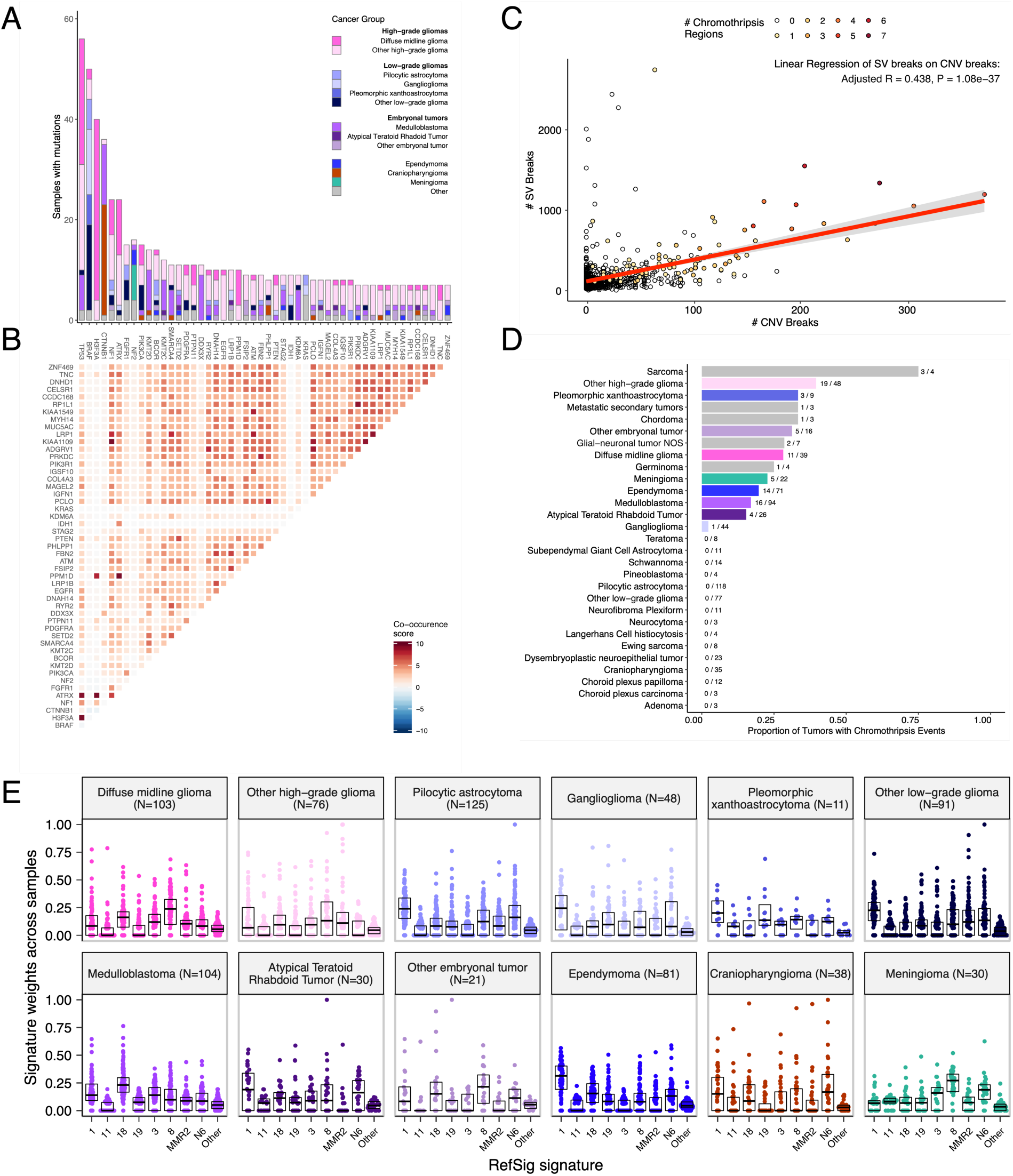
Mutational co-occurrence and signatures highlight key oncogenic drivers. A, Bar plot of occurrence and co-occurrence of nonsynonymous mutations for the 50 most commonly mutated genes across all tumor types (annotated from `cancer_group` if N >= 10 or `Other` if N < 10); B, Co-occurrence and mutual exclusivity of nonsynonymous mutations between genes; The co-occurrence score is defined as *I*(–log_10_(*P*)) where *P* is defined by Fisher’s exact test and *I* is 1 when mutations co-occur more often than expected and −1 when exclusivity is more common; C, The number of SV breaks significantly correlates with the number of CNV breaks (Adjusted R = 0.438, p = 1.08e-37). D, Chromothripsis frequency across pediatric brain tumors shown by `cancer_group` with N >= 3. E, Sina plots of RefSig signature weights for signatures 1, 11, 18, 19, 3, 8, N6, MMR2, and Other across cancer groups. Box plot lines represent the first quartile, median, and third quartile.

### Transcriptomic Landscape of Pediatric Brain Tumors

#### Prediction of *TP53* oncogenicity and telomerase activity

To understand the *TP53* phenotype in each tumor, we ran a classifier previously trained on TCGA^38^ to calculate a *TP53* score and infer *TP53* inactivation status. We compared results of this classifier to “true positive” alterations derived using high-confidence SNVs, CNVs, SVs, and fusions in *TP53*. Specifically, we annotated *TP53* alterations as “activated” if samples harbored one of p.R273C or p.R248W gain-of-function mutations^39^, or “lost” if the given patient either had a Li Fraumeni Syndrome (LFS) predisposition diagnosis, the tumor harbored a known hotspot mutation, or the tumor contained two hits (e.g. both SNV and CNV), which would suggest both alleles had been affected. If the *TP53* mutation did not reside within the DNA-binding domain or we did not detect any alteration in *TP53*, we annotate the tumor as “other,” reflecting its unknown *TP53* alteration status. The classifier achieved a high accuracy (AUROC = 0.85) for rRNA-depleted, stranded samples compared to randomly shuffled *TP53* scores (Figure 4A). By contrast, while this classifier has previously shown strong performance on poly-A data from both adult^38^ tumors and pediatric patient-derived xenografts^40^, it did not perform as well on the poly-A samples in this cohort (AUROC = 0.62; Figure S5A).

While we expected that samples annotated as “lost” would have higher *TP53* scores than would samples annotated as “other,” we observed that samples annotated as “activated” had similar *TP53* scores to those annotated as “lost” (Figure 4B, Wilcoxon p = 0.23). This result suggests that the classifier actually detects an oncogenic, or altered, *TP53* phenotype (scores > 0.5) rather than solely *TP53* inactivation, as interpreted previously^38^. Moreover, tumors with “activating” *TP53* mutations showed higher *TP53* expression compared to those with *TP53* “loss” mutations (Wilcoxon p = 3.5e-3, Figure 4C). Tumor types with the highest median *TP53* scores were those known to harbor somatic *TP53* alterations and included DMGs, medulloblastomas, HGGs, DNETs, ependymomas, and craniopharyngiomas (Figure 4D), while gangliogliomas, LGGs, meningiomas, and schwannomas had the lowest median scores.

To further validate the classifier’s accuracy, we assessed *TP53* scores for patients with LFS, hypothesizing that all of these tumors would have high scores. Indeed, we observed higher scores in LFS tumors (N = 8) for which we detected high-confidence *TP53* somatic alterations (**Tables S1 and S3**). Although we did not detect canonical somatic *TP53* mutations in two patients whose tumors had low *TP53* scores (BS_DEHJF4C7 with a score of 0.09 and BS_ZD5HN296 with a score of 0.28), we confirmed from pathology reports these patients were both diagnosed with LFS and had pathogenic germline variants in *TP53*. In addition, the tumor purity of these two LFS samples was low (16% and 37%, respectively), suggesting the classifier may require a certain level of tumor purity to achieve good performance, as we expect *TP53* to be intact in normal cells.

We next used gene expression data to predict telomerase activity using EXpression-based Telomerase ENzymatic activity Detection (`EXTEND`)^41^ as a surrogate measure of malignant potential^41, 42^, such that higher `EXTEND` scores suggest increased malignant potential. As expected, `EXTEND` scores significantly correlated with *TERC* (R = 0.619, p < 0.01) and *TERT* (R = 0.491, p < 0.01) expression (Figure S5B-C). We found aggressive tumors such as HGGs (DMGs and other high-grade gliomas) and MB had high `EXTEND` scores (Figure 4D), while low-grade lesions such as schwannomas, GNGs, DNETs, and other low-grade gliomas had among the lowest scores (**Table S3**). These findings support previous reports of a more aggressive phenotype in tumors with higher telomerase activity^43–46^.

#### Hypermutant tumors share mutational signatures and have dysregulated ***TP53***

We further investigated the mutational signature profiles of the hypermutant (TMB > 10 Mut/Mb; N = 3) and ultra-hypermutant (TMB > 100 Mut/Mb; N = 4) tumors and/or derived cell lines from six patients in the OpenPBTA cohort (Figure 4E). Five of six tumors were diagnosed as HGGs and one was a brain metastasis of a MYCN non-amplified neuroblastoma tumor. Signature 11, which is associated with exposure to temozolomide plus *MGMT* promoter and/or mismatch repair deficiency^47^, was indeed present in tumors with previous exposure to the drug (Table 2). We detected the MMR2 signature in tumors of four patients (PT_0SPKM4S8, PT_3CHB9PK5, PT_JNEV57VK, and PT_VTM2STE3) diagnosed with either constitutional mismatch repair deficiency (CMMRD) or Lynch syndrome (Table 2), genetic predisposition syndromes caused by a variant in a mismatch repair gene such as *PMS2*, *MLH1*, *MSH2*, *MSH6*, or others^48^. Three of these patients harbored pathogenic germline variants in one of the aforementioned genes. While we did not find a *known* pathogenic variant in the germline of PT_VTM2STE3, this patient had a self-reported *PMS2* variant noted in their pathology report and we did find 19 intronic variants of unknown significance (VUS) in *PMS2*. This is not surprising since an estimated 49% of germline *PMS2* variants in patients with CMMRD and/or Lynch syndrome are VUS^48^. Interestingly, while the cell line derived from patient PT_VTM2STE3’s tumor at progression was not hypermutated (TMB = 5.7 Mut/Mb), it solely showed the MMR2 signature of the eight CNS signatures examined, suggesting selective pressure to maintain a mismatch repair (MMR) phenotype *in vitro*. From patient PT_JNEV57VK, only one of the two cell lines derived from the progressive tumor was hypermutated (TMB = 35.9 Mut/Mb). This hypermutated cell line was strongly weighted towards signature 11, while this patient’s non-hypermutated cell line showed a number of lesser signature weights (1, 11, 18, 19, MMR2; Table S2), highlighting the plasticity of mutational processes and the need to carefully genomically characterize and select models for preclinical studies based on research objectives.

We observed that signature 18, which has been associated with high genomic instability and can lead to a hypermutator phenotype^36^, was uniformly represented among hypermutant solid tumors. Additionally, we found that all of the HGG tumors or cell lines had dysfunctional *TP53* (Table 2), consistent with a previous report showing *TP53* dysregulation is a dependency in tumors with high genomic instability^36^. With one exception, hypermutant and ultra-hypermutant tumors had high *TP53* scores (> 0.5) and telomerase activity. Interestingly, none of the hypermutant samples showed evidence of signature 3 (present in homologous recombination deficient tumors), signature 8 (arises from double nucleotide substitutions/unknown etiology), or signature N6 (a universal CNS tumor signature). The mutual exclusivity of signatures 3 and MMR2 corroborates a previous report suggesting tumors do not tend to feature both deficient homologous repair and mismatch repair^38^.

**Table 2.** Patients with hypermutant tumors. Listed are patients with at least one hypermutant or ultra-hypermutant tumor or cell line. Pathogenic (P) or likely pathogenic (LP) germline variants, coding region TMB, phase of therapy, therapeutic interventions, cancer predisposition (CMMRD = Constitutional mismatch repair deficiency), and molecular subtypes are included.

| Kids First Participant ID | Kids First Biospecimen ID | CBTN ID | Phase of therapy | Composition | Therapy post-biopsy | Cancer predisposition | Pathogenic germline variant | TMB | OpenPBTA molecular subtype |
| --- | --- | --- | --- | --- | --- | --- | --- | --- | --- |
| PT_0SPKM4S8 | BS_VW4XN9Y7 | 7316-2640 | Initial CNS Tumor | Solid Tissue | Radiation, Temozolomide, CCNU | None documented | NM_000535.7(PMS2):c.137G>T (p.Ser46Ile) (LP) | 187.4 | HGG, H3 wildtype, TP53 activated |
| PT_3CHB9PK5 | BS_20TBZG09 | 7316-515 | Initial CNS Tumor | Solid Tissue | Radiation, Temozolomide, Irinotecan, Bevacizumab | CMMRD | NM_000179.3(MSH6):c.3439-2A>G (LP) | 307 | HGG, H3 wildtype, TP53 loss |
| PT_3CHB9PK5 | BS_8AY2GM4G | 7316-2085 | Progressive | Solid Tissue | Radiation, Temozolomide, Irinotecan, Bevacizumab | CMMRD | NM_000179.3(MSH6):c.3439-2A>G (LP) | 321.6 | HGG, H3 wildtype, TP53 loss |
| PT_EB0D3BXG | BS_F0GNWEJJ | 7316-3311 | Progressive | Solid Tissue | Radiation, Nivolumab | None documented | None detected | 26.3 | Metastatic NBL, MYCN non-amplified |
| PT_JNEV57VK | BS_85Q5P8GF | 7316-2594 | Initial CNS Tumor | Solid Tissue | Radiation, Temozolomide | Lynch Syndrome | NM_000251.3(MSH2):c.1906G>C (p.Ala636Pro) (P) | 4.7 | DMG, H3 K28, TP53 loss |
| PT_JNEV57VK | BS_HM5GFJN8 | 7316-3058 | Progressive | Derived Cell Line | Radiation, Temozolomide, Nivolumab | Lynch Syndrome | NM_000251.3(MSH2):c.1906G>C (p.Ala636Pro) (P) | 35.9 | DMG, H3 K28, TP53 loss |
| PT_JNEV57VK | BS_QWM9BPDY | 7316-3058 | Progressive | Derived Cell Line | Radiation, Temozolomide, Nivolumab | Lynch Syndrome | NM_000251.3(MSH2):c.1906G>C (p.Ala636Pro) (P) | 7.4 | DMG, H3 K28, TP53 loss |
| PT_JNEV57VK | BS_P0QJ1QAH | 7316-3058 | Progressive | Solid Tissue | Radiation, Temozolomide, Nivolumab | Lynch Syndrome | NM_000251.3(MSH2):c.1906G>C (p.Ala636Pro) (P) | 6.3 | DMG, H3 K28, TP53 activated |
| PT_S0Q27J13 | BS_P3PF53V8 | 7316-2307 | Initial CNS Tumor | Solid Tissue | Radiation, Temozolomide, Irinotecan | None documented | None detected | 15.5 | HGG, H3 wildtype, TP53 activated |
| PT_VTM2STE3 | BS_ERFMPQN3 | 7316-2189 | Progressive | Derived Cell Line | Unknown | Lynch Syndrome | None detected | 5.7 | HGG, H3 wildtype, TP53 loss |
| PT_VTM2STE3 | BS_02YBZSBY | 7316-2189 | Progressive | Solid Tissue | Unknown | Lynch Syndrome | None detected | 274.5 | HGG, H3 wildtype, TP53 activated |

Next, we asked whether transcriptomic classification of *TP53* dysregulation and/or telomerase activity recapitulate the known prognostic influence of these oncogenic biomarkers. To this end, we conducted a multivariate Cox regression on overall survival (Figure 4F; **STAR Methods**), controlling for extent of tumor resection and whether a tumor was low-grade (LGG group) or high-grade (HGG group). We identified several expected trends, including a significant overall survival benefit if the tumor had been fully resected (HR = 0.35, 95% CI = 0.2 - 0.62, p < 0.001) or if the tumor belonged to the LGG group (HR = 0.046, 95% CI = 0.0062 - 0.34, p = 0.003) as well as a significant risk if the tumor belonged to the HGG group (HR = 6.2, 95% CI = 4.0 - 9.5, p < 0.001). High telomerase scores were associated with poor prognosis across brain tumor histologies (HR = 20, 95% CI = 6.4 - 62, p < 0.001), demonstrating that `EXTEND` scores calculated from RNA-Seq are an effective rapid surrogate measure for telomerase activity. Although higher *TP53* scores, which predict *TP53* gene or pathway dysregulation, were not a significant predictor of risk across the entire OpenPBTA cohort (**Table S4**), we did find a significant survival risk associated with higher *TP53* scores within DMGs (HR = 6436, 95% CI = 2.67 - 1.55e7, p = 0.03) and ependymomas (HR = 2003, 95% CI = 9.9 - 4.05e5, p = 0.005). Since we observed the negative prognostic effect of *TP53* scores for HGGs, we assessed the effect of molecular subtypes within HGG samples on survival risk. We found that DMG H3 K28 tumors with *TP53* loss had significantly worse prognosis (HR = 2.8, CI = 1.4-5.6, p = 0.003) than did DMG H3 K28 tumors with wildtype *TP53* (Figure 4G and Figure 4H). This finding was also recently reported in a restrospective analysis of DIPG tumors from the PNOC003 clinical trial^12^.

**Figure 4.**
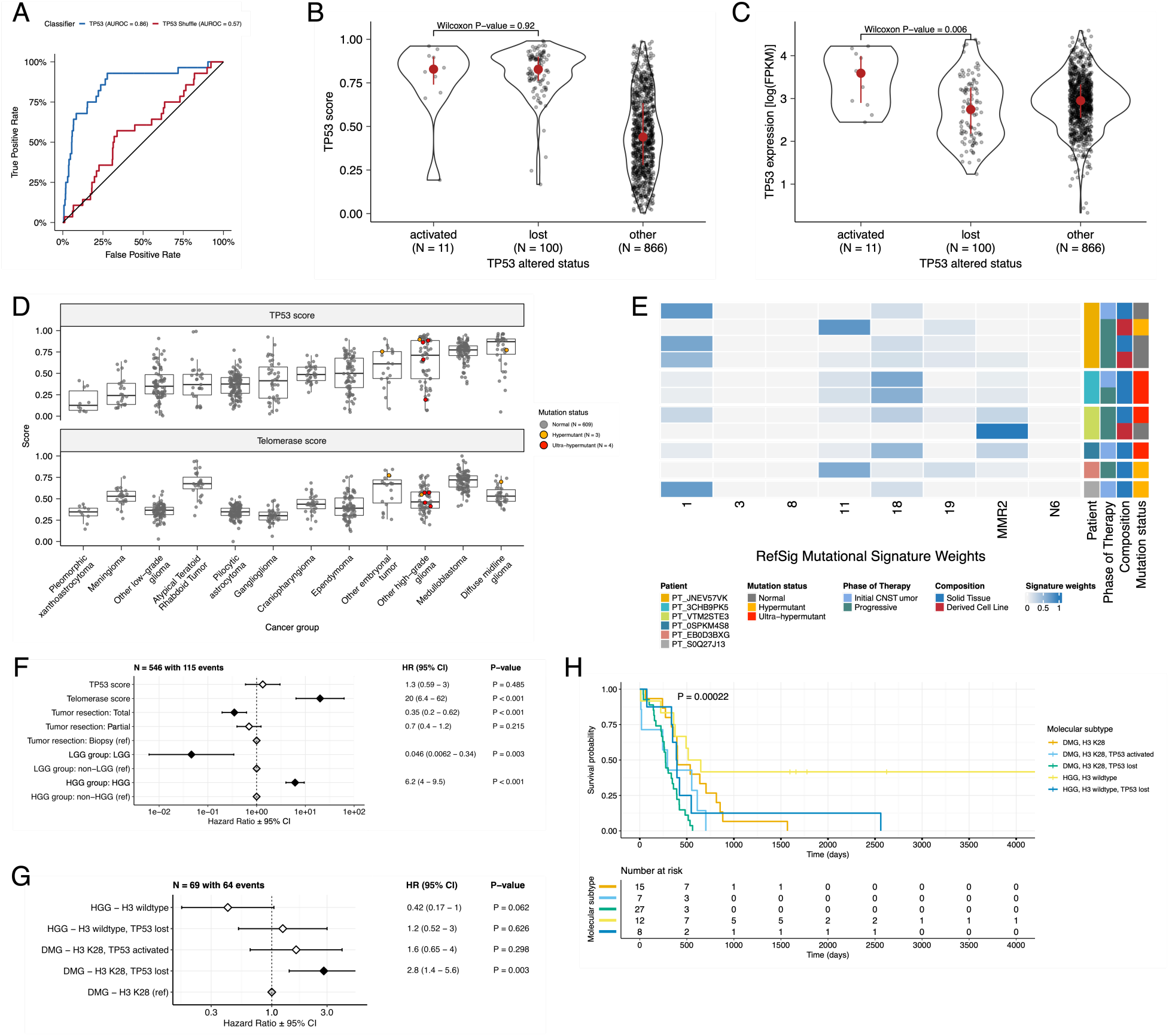
*TP53* and telomerase activity. A, Receiver Operating Characteristic for *TP53* classifier run on FPKM of stranded RNA-Seq samples. B, Violin and box plots of *TP53* scores plotted by *TP53* alteration type (Nactivated = 11, Nlost = 100, Nother = 866). C, Violin and box plots of *TP53* RNA expression plotted by *TP53* activation status (Nactivated = 11, Nlost = 100, Nother = 866). D, Box plots of *TP53* and telomerase (EXTEND) scores grouped by `cancer_group`. Mutation status is highlighted in orange (hypermutant) or red (ultra-hypermutant). E, Heatmap of RefSig mutational signatures for patients who have least one tumor or cell line with a TMB >= 10 Mut/Mb. F, Forest plot depicting the prognostic effects of *TP53* and telomerase scores on overall survival, controlling for extent of tumor resection, LGG group, and HGG group. G, Forest plot depicting the effect of molecular subtype on overall survival of HGGs. For F and G, hazard ratios (HR) with 95% confidence intervals and p-values are listed. Significant p-values are denoted with black diamonds. Reference groups are denoted by grey diamonds. H, Kaplan-Meier curve of HGG tumors by molecular subtype.

#### Histologic and oncogenic pathway clustering

UMAP visualization of gene expression variation across brain tumors (Figure 5A) showed the expected clustering of brain tumors by histology. We additionally explored UMAP projections of gene expression within molecular subtypes for certain cancer groups. We observed that, except for three outliers, *C11orf95::RELA* (*ZFTA::RELA*) fusion-positive ependymomas fell within distinct clusters (Figure S6A). Medulloblastoma (MB) samples cluster by molecular subtype, with WNT and SHH in distinct clusters and Groups 3 and 4 showing some overlap (Figure S6B), as expected. Of note, two MB samples annotated as the SHH subtype did not cluster with the other MB samples, and one clustered with Group 3 and 4 samples, suggesting potential subtype misclassification or different underlying biology of these two tumors. *BRAF*-driven low-grade gliomas (Figure S6C) were present in three separate clusters, suggesting that there might be additional shared biology within each cluster. Histone H3 G35-mutant high-grade gliomas generally clustered together and away from K28-mutant tumors (Figure S6D). Interestingly, although H3 K28-mutant tumors have different biological drivers than do H3 wildtype tumors^49^, they did not form distinct clusters. This pattern suggests these subtypes may be driven by common transcriptional programs, have other much stronger biological drivers than their known distinct epigenetic drivers, or our sample size is too small to detect transcriptional differences.

We next performed gene set variant analysis (GSVA) for Hallmark cancer gene sets to demonstrate activation of underlying oncogenic pathways (Figure 5B and quantified immune cell fractions across OpenPBTA tumors using quanTIseq (Figure 5C and Figure S6E). Through these analyses, we were able to recapitulate previously-described tumor biology. For example, HGG, DMG, MB, and ATRT tumors are known to upregulate *MYC*^50^ which in turn activates *E2F* and S phase^51^. Indeed, we detected significant (Bonferroni-corrected p < 0.05) upregulation of *MYC* and *E2F* targets, as well as G2M (cell cycle phase following S phase) in MBs, ATRTs, and HGGs compared to several other cancer groups. In contrast, LGGs showed significant downregulation (Bonferroni-corrected p < 0.05) of these pathways. Schwannomas and neurofibromas, which have a documented inflammatory immune microenvironment of T and B lymphocytes as well as tumor-associated macrophages (TAMs), are driven by upregulation of cytokines such as IFNγ, IL-1, and IL-6, and TNFα^52^. Indeed, we observed significant upregulation of these cytokines in GSVA hallmark pathways (Bonferroni-corrected p < 0.05) (Figure 5B) and found immune cell types dominated by monocytes in these tumors (Figure 5C). We also observed significant upregulation of pro-inflammatory cytokines IFNα and IFNγ in LGGs and craniopharyngiomas compared to medulloblastoma and ependymoma tumors (Bonferroni-corrected p < 0.05), both of which showed significant down-regulation of these cytokines (Figure 5B). Together, these results supported previous proteogenomic findings of lower immune infiltration in aggressive medulloblastomas and ependymomas versus higher immune infiltration in *BRAF*-driven LGG and craniopharyngiomas^53^.

Although CD8+ T-cell infiltration across all cancer groups was quite low (Figure 5C), we observed some signal in specific cancer molecular subtypes (Groups 3 and 4 medulloblastoma) as well as outlier tumors (BRAF-driven LGG, BRAF-driven and wildtype ganglioglioma, and CNS embryonal NOS; Figure S6E) Surprisingly, the classically immunologically-cold HGG and DMG tumors^54, 55^ contained higher overall fractions of immune cells, where monocytes, dendritic cells, and NK cells were the most prevalent (Figure 5C). Thus, we suspect that quanTIseq might actually have captured microglia within these immune cell fractions.

While we did not detect notable prognostic effects of immune cell infiltration on overall survival in HGG or DMG tumors, we did find that high levels of macrophage M1 and monocytes were associated with poorer overall survival (monocyte HR = 2.1e18, 95% CI = 3.80e5 - 1.2e31, p = 0.005) in medulloblastoma tumors (Figure 5D). We further reproduced previous findings (Figure 5E) that medulloblastomas typically have low expression of *CD274* (PD-L1)^56^. However, we also found that higher expression of *CD274* was significantly associated with improved overall prognosis for medulloblastoma samples, although with a marginal effect size (HR = 0.0012, 95% CI = 7.5e−06 - 0.18, p = 0.008) (Figure 5D). This result may be explained by the higher expression of *CD274* found in WNT subtype tumors by us and others^57^, as this diagnosis carries the best prognosis of all medulloblastoma subgroups (Figure 5E).

Finally, we asked whether any molecular subtypes might show an immunologically-hot phenotype, as roughly defined by a greater proportion of CD8+ to CD4+ T cells^58, 59^. While adamantinomatous craniopharyngiomas and Group 3 and Group 4 medulloblastomas had the highest CD8+ to CD4+ T cell ratios (Figure S6F), very few tumors had ratios greater than 1, highlighting an urgent need to identify novel therapeutics for these immunologically-cold pediatric brain tumors with poor prognosis.

**Figure 5.**
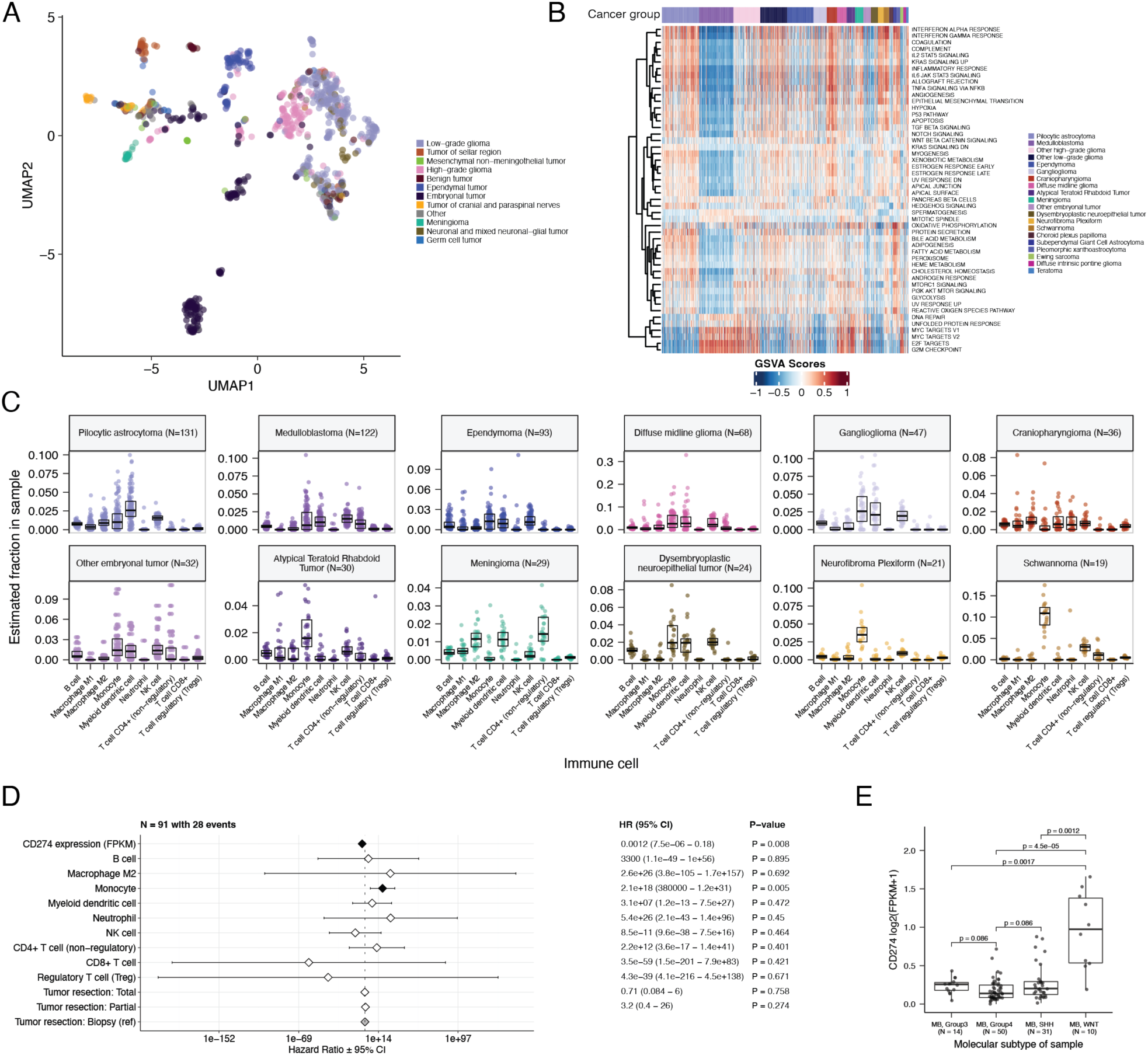
Transcriptomic and immune landscape of pediatric brain tumors. A, First two dimensions from UMAP of sample transcriptome data. Points are colored by the broad histology of the samples they represent. B, Heatmap of GSVA scores for Hallmark gene sets with significant differences, with samples ordered by cancer group. C, Box plots of quanTIseq estimates of immune cell proportions in select cancer groups with N > 15 samples. Note: Other HGGs and other LGGs have immune cell proportions similar to DMG and pilocytic astrocytoma, respectively, and are not shown. D, Forest plot depicting the additive effects of *CD274* expression, immune cell proportion, and extent of tumor resection on overall survival of medulloblastoma patients. Hazard ratios (HR) with 95% confidence intervals and p-values are listed. Significant p-values are denoted with black diamonds. Reference groups are denoted by grey diamonds. Of note, the Macrophage M1 HR was 0 (coefficient = −9.90e+4) with infinite upper and lower CIs, and thus it was not included in the figure. E, Box plot of *CD274* expression (log2 FPKM) for medulloblastoma samples grouped by molecular subtype. Bonferroni-corrected p-values from Wilcoxon tests are shown.

## Discussion

We created OpenPBTA to define an open, real-time, reproducible analysis framework to genomically characterize pediatric brain tumors that brings together basic and translational researchers, clinicians, and data scientists on behalf of accelerated discovery and clinical impact. We provide robust reusable code and data resources, paired with cloud-based availability of source and derived data resources, to the pediatric oncology community, encouraging interdisciplinary scientists to collaborate on new analyses in order to accelerate therapeutic translation for children with cancer, goals we are seeing play out in real-time. To our knowledge, this initiative represents the first large-scale, collaborative, open analysis of genomic data coupled with open manuscript writing, in which we comprehensively analyzed the largest cohort of pediatric brain tumors to date, comprising 1,074 tumors across 58 distinct histologies. We used available WGS, WXS, and RNA-Seq data to generate high-confidence consensus SNV and CNV calls, prioritize putative oncogenic fusions, and establish over 40 scalable modules to perform common downstream cancer genomics analyses, all of which have undergone rigorous scientific and analytical code review. We detected and showed expected patterns of genomic lesions, mutational signatures, and aberrantly regulated signaling pathways across multiple pediatric brain tumor histologies.

Assembling large, pan-histology cohorts of fresh frozen samples and associated clinical phenotypes and outcomes requires a multi-year, multi-institutional framework, like those provided by CBTN and PNOC. As such, uniform clinical molecular subtyping was largely not performed for most of this cohort at the time of diagnosis and/or at surgery, and when available (e.g., sparse medulloblastoma subtypes), it required manual curation from pathology reports and/or free text clinical data fields. Furthermore, rapid classification to derive molecular subtypes could not be immediately performed since research-based DNA methylation data for these samples are not yet available. Thus, to enable biological interrogation of specific tumor subtypes, we created RNA- and DNA-based subtyping modules aligned with WHO molecularly-defined diagnoses. We worked closely with pathologists and clinicians to build modules from which we determined a research-grade integrated diagnosis for 60% of samples while discovering incorrectly diagnosed or mis-identified samples in the OpenPBTA cohort.

We harnessed RNA expression data for a number of analyses, yielding important biological insights across multiple brain tumor histologies. For example, we performed subtyping of medulloblastoma tumors, for which only 35% (43/122) had subtype information from pathology reports. Among the subtyped tumors, we accurately recapitulated subtypes using MM2S (91%; 39/43) or medulloPackage (95%; 41/43)^60, 61^. We then applied the consensus of these methods to subtype all medulloblastoma tumors lacking pathology-based subtypes.

We advanced the integrative analyses and cross-cohort comparison via a number of validated modules. We used an expression classifier to determine whether tumors have dysfunctional *TP53*^38^ and the EXTEND algorithm to determine their degree of telomerase activity using a 13-gene signature^41^. Interestingly, in contrast to adult colorectal cancer and gastric adenocarcinoma, in which *TP53* loss of function is less frequent in hypermutated tumors^62, 63^, we found that hypermutant HGG tumors universally displayed dysregulation of *TP53*. Furthermore, high *TP53* scores were a significant prognostic marker for poor overall survival for patients with certain tumor types, such as H3 K28-altered DMGs and ependymomas. We also show that EXTEND scores are a robust surrogate measure for telomerase activity in pediatric brain tumors. By assessing *TP53* and telomerase activity prospectively from expression data, information usually only attainable with DNA sequencing and/or qPCR, we can quickly incorporate oncogenic biomarker and prognostic knowledge and expand our biological understanding of these tumors.

We identified enrichment of hallmark cancer pathways and characterized the immune cell landscape across pediatric brain tumors, demonstrating tumors in some histologies, such as schwannomas, craniopharyngiomas, and low-grade gliomas, may have a inflammatory tumor microenvironment. Of note, we observed upregulation of IFNγ, IL-1, and IL-6, and TNFα in craniopharyngiomas, tumors difficult to resect due to their anatomical location and critical surrounding structures. Neurotoxic side effects have been reported when interferon alpha immunotherapy is administered to reduce cystic craniopharyngioma tumor size and/or delay progression^64, 65^. Thus, additional immune vulnerabilities, such as IL-6 inhibition and immune checkpoint blockade, have recently been proposed as therapies for cystic adamantinomatous craniopharyngiomas^66–70^ and our results noted above support this approach. Finally, our study reproduced the overall known poor infiltration of CD8+ T cells and general low expression of *CD274* (PD-L1) in pediatric brain tumors, further highlighting the urgent need to identify novel therapeutic strategies for these immunologically cold tumors.

OpenPBTA has rapidly become a foundational data analysis and processing layer for a number of discovery research and translational projects which will continue to add other genomic modalities and analyses, such as germline, methylation, single cell, epigenomic, mRNA splicing, imaging, and model drug response data. For example, the RNA fusion filtering module created within OpenPBTA set the stage for development of the R package *annoFuse*^71^ and an R Shiny application *<u>shinyFuse</u>*. Using medulloblastoma subtyping and immune deconvolution analyses performed herein, Dang and colleagues showed enrichment of monocyte and microglia-derived macrophages within the SHH subgroup which they suggest may accumulate following radiation therapy^10^. Expression and copy number analyses were used to demonstrate that *GPC2* is a highly expressed and copy number gained immunotherapeutic target in ETMRs, medulloblastomas, choroid plexus carcinomas, H3 wildtype high-grade gliomas, as well as DMGs. This led Foster and colleagues to subsequently develop a chimeric antigen receptor (CAR) directed against *GPC2*, for which they show preclinical efficacy in mouse models^11^. Moreover, OpenPBTA has enabled a framework to support real-time integration of clinical trial subjects as each was enrolled on the PNOC008 high-grade glioma clinical trial^72^, allowing researchers and clinicians to link tumor biology to translational impact through clinical decision support during tumor board discussions. Finally, as part of the the NCI’s Childhood Cancer Data Initiative (CCDI), the OpenPBTA project was recently expanded into a pan-pediatric cancer effort (https://github.com/PediatricOpenTargets/OpenPedCan-analysis) to build the Molecular Targets Platform (https://moleculartargets.ccdi.cancer.gov/) in support of the RACE Act. An additional, large-scale cohort of >2,500 tumor samples and associated germline DNA is in the process of undergoing sequence data generation as part of CBTN CCDI-Kids First NCI and Common Fund project (https://commonfund.nih.gov/kidsfirst/2021X01projects#FY21_Resnick). Like the original OpenPBTA cohort, data will be processed and released in near real-time via the Kids First Data Resource and integrated with OpenPBTA. The OpenPBTA project has paved the way for new modes of collaborative data-driven discovery, open, reproducible, and scalable analyses that will extend beyond the current research described herein, and we anticipate this foundational work will continue to have a long-term impact within the pediatric brain tumor translational research community and beyond, ultimately leading to accelerated impact and improved outcomes for children with cancer.

All code and processed data are openly available through GitHub, CAVATICA, and PedcBioPortal (see **STAR METHODS**).

## Acknowledgments

We graciously thank the patients and families who have donated their tumors to the Children’s Brain Tumor Network (CBTN) and/or the Pacific Pediatric Neuro-oncology Consortium, without which this research would not be possible.

Philanthropic support has ensured the CBTN’s ability to collect, store, manage, and distribute specimen and data since it was founded in 2013. In addition to the support from the CBTN Executive Council members and Brain Tumor Board of Visitors, the following donors have provided leadership level support for CBTN: Children’s Brain Tumor Foundation, Easie Family Foundation, Kortney Rose Foundation, Lilabean Foundation, Minnick Family Charitable Fund, Perricelli Family, Psalm 103 Foundation, and Swifty Foundation.

This work was funded through the Alex’s Lemonade Stand Foundation (ALSF) Childhood Cancer Data Lab (CSG), ALSF Young Investigator Award (JLR), ALSF Catalyst Award (JLR, ACR, PBS), ALSF Catalyst Award (SJS), ALSF CCDL Postdoctoral Training Grant (SMF), Children’s Hospital of Philadelphia Division of Neurosurgery (PBS and ACR), the Australian Government, Department of Education (APH), the St. Anna Kinderkrebsforschung, Austria (ARP), the Mildred Scheel Early Career Center Dresden P2, funded by the German Cancer Aid (ARP), and NIH Grants 3P30 CA016520-44S5 (ACR), U2C HL138346-03 (ACR, APH), U24 CA220457-03 (ACR), K12GM081259 (SMF), R03-CA23036 (SJD), and NIH Contract No. HHSN261200800001E (SJD). This project has been funded in part with Federal funds from the National Cancer Institute, National Institutes of Health, under Contract No. 75N91019D00024, Task Order No. 75N91020F00003 (JLR, ACR, APH). Additionally, this work was supported by the Intramural Research Program of the Division of Cancer Epidemiology and Genetics of the National Cancer Institute. The content of this publication does not necessarily reflect the views or policies of the Department of Health and Human Services, nor does mention of trade names, commercial products or organizations imply endorsement by the U.S. Government.

The authors would like to thank the following collaborators who contributed or supervised analyses present in the analysis repository that were not included in the manuscript: William Amadio, Holly C. Beale, Ellen T. Kephart, A. Geoffrey Lyle, and Olena M. Vaske. Finally, we would like to thank Yuanchao Zhang and Eric Wafula for adding to the project codebase, Jessica B. Foster for helpful discussions while drafting the manuscript, and Gina D. Mawla for identifying and reporting issues in OpenPBTA data.

## Author Contributions

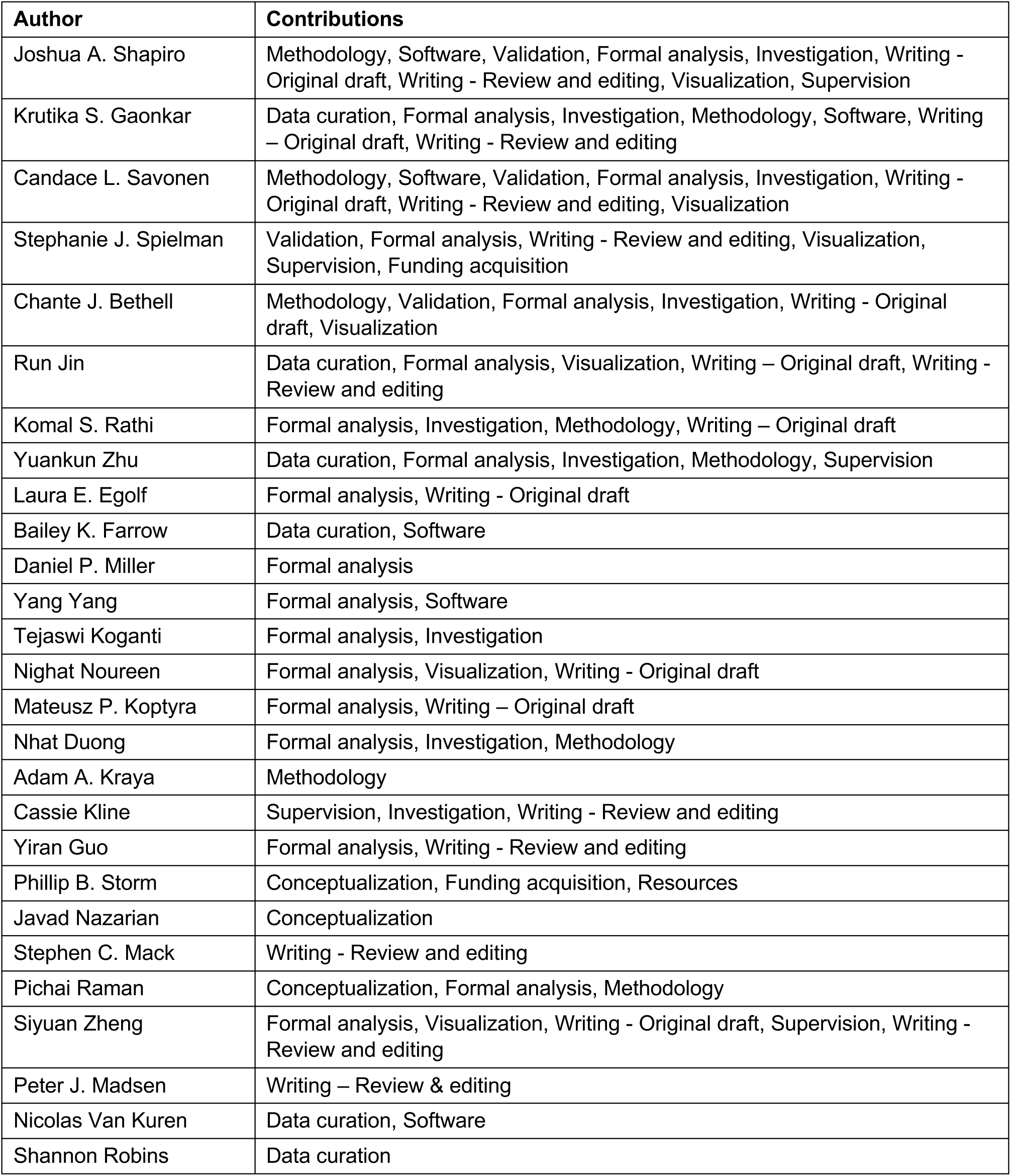
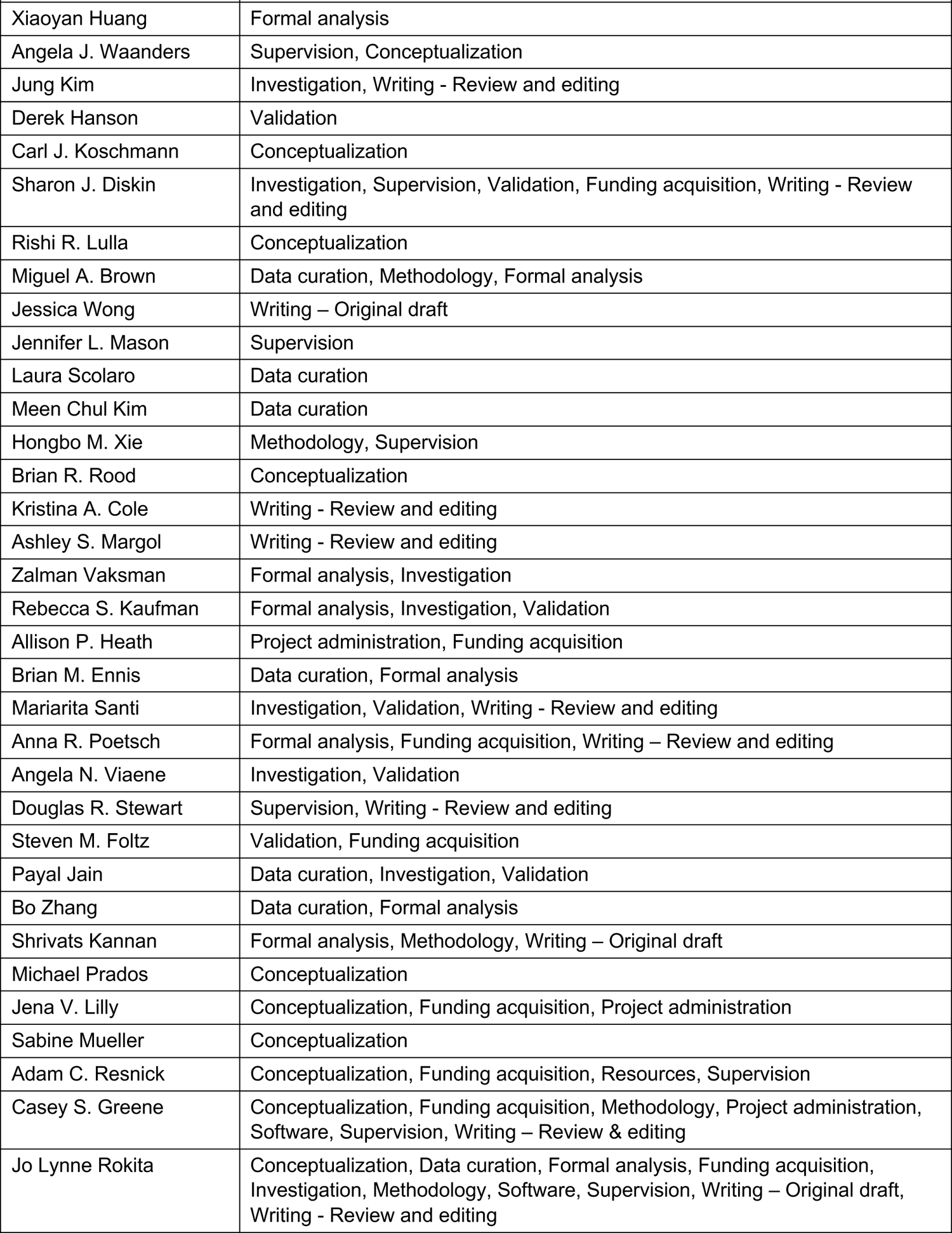
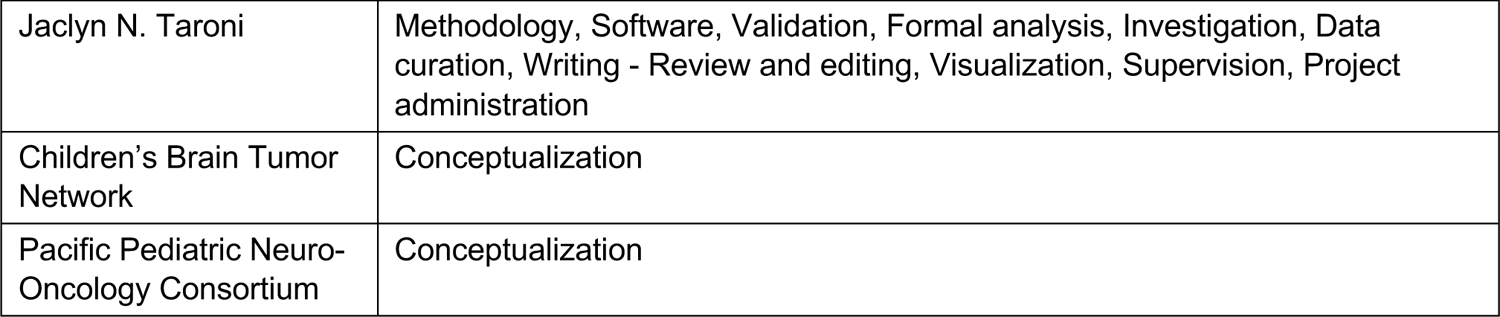

Except for the first and last four authors, authorship order was determined as follows: Authors who contributed to the OpenPBTA code base are listed based on number of modules included in the manuscript to which that individual contributed and, in the case of ties, a random order is used. All remaining authors are then listed in a random order.

Code for determining authorship order can be found in the `count-contributions` module of the OpenPBTA analysis repository.

## STAR METHODS

### RESOURCE AVAILABILITY

#### Lead contact

Requests for access to OpenPBTA raw data and/or specimens may be directed to, and will be fulfilled by Jo Lynne Rokita.

#### Materials availability

This study did not create new, unique reagents.

#### Data and code availability

Raw and harmonized WGS, WXS, and RNA-Seq data derived from human samples are available within the KidsFirst Portal^73^ upon access request to the CBTN (https://cbtn.org/) as of the date of the publication. In addition, merged summary files are openly accessible at https://cavatica.sbgenomics.com/u/cavatica/openpbta or via download script in the https://github.com/AlexsLemonade/OpenPBTA-analysis repository. Summary data are visible within PedcBioPortal at https://pedcbioportal.kidsfirstdrc.org/study/summary?id=openpbta. Associated DOIs are listed in the **Key Resources Table**.

All original code was developed within the following repositories and is publicly available as follows. Primary data analyses can be found at https://github.com/d3b-center/OpenPBTA-workflows. Downstream data analyses can be found at https://github.com/AlexsLemonade/OpenPBTA-analysis. Manuscript code can be found at https://github.com/AlexsLemonade/OpenPBTA-manuscript. Associated DOIs are listed in the **Key Resources Table**. Software versions are documented in Table S5 as an appendix to the **Key Resources Table**.

Any additional information required to reanalyze the data reported in this paper is available from the lead contact upon request.

### METHOD DETAILS

#### Biospecimen Collection

The Pediatric Brain Tumor Atlas specimens are comprised of samples from Children’s Brain Tumor Network (CBTN) and the Pediatric Pacific Neuro-Oncology Consortium (PNOC). The <u>CBTN</u> is a collaborative, multi-institutional (26 institutions worldwide) research program dedicated to the study of childhood brain tumors. <u>PNOC</u> is an international consortium dedicated to bringing new therapies to children and young adults with brain tumors. We also include blood and tumor biospecimens from newly-diagnosed diffuse intrinsic pontine glioma (DIPG) patients as part of the PNOC003 clinical trial PNOC003/NCT02274987^15^.

The CBTN-generated cell lines were derived from either fresh tumor tissue directly obtained from surgery performed at Children’s Hospital of Philadelphia (CHOP) or from prospectively collected tumor specimens stored in Recover Cell Culture Freezing medium (cat# 12648010, Gibco). We dissociated tumor tissue using enzymatic method with papain as described^14^. Briefly, we washed tissue with HBSS (cat# 14175095, Gibco), and we minced and incubated the tissue with activated papain solution (cat# LS003124, SciQuest) for up to 45 minutes. We used ovomucoid solution (cat# 542000, SciQuest) to inactivate the papain, briefly treated tissue with DNase (cat# 10104159001, Roche), passed it through the 100μm cell strainer (cat# 542000, Greiner Bio-One). We initiated two cell culture conditions based on the number of cells available. For cultures utilizing the fetal bovine serum (FBS), we plated a minimum density of 3×105 cells/mL in DMEM/F-12 medium (cat# D8062, Sigma) supplemented with 20% FBS (cat# SH30910.03, Hyclone), 1% GlutaMAX (cat# 35050061, Gibco), Penicillin/Streptomycin-Amphotericin B Mixture (cat# 17-745E, Lonza), and 0.2% Normocin (cat# ant-nr-2, Invivogen). For serum-free media conditions, we plated cells at minimum density of 1×106 cells/mL in DMEM/F12 medium supplemented with 1% GlutaMAX, 1X B-27 supplement minus vitamin A (cat# 12587-010, Gibco), 1x N-2 supplement (cat# 17502001, Gibco), 20 ng/ml epidermal growth factor (cat# PHG0311L, Gibco), 20 ng/mL basic fibroblast growth factor (cat# 100-18B, PeproTech), 2.5μg/mL heparin (cat# H3149, Sigma), Penicillin/Streptomycin-Amphotericin B Mixture, and 0.2% Normocin.

### Nucleic acids extraction and library preparation

#### PNOC samples

The Translational Genomic Research Institute (TGEN; Phoenix, AZ) performed DNA and RNA extractions on tumor biopsies using a DNA/RNA AllPrep Kit (Qiagen, #80204). All RNA used for library prep had a minimum RIN of seven, but no QC thresholds were implemented for the DNA. For library preparation, 500 ng of nucleic acids were used as input for RNA-Seq, WXS, and targeted DNA panel (panel) sequencing. RNA library preparation was performed using the TruSeq RNA Sample Prep Kit (Illumina, #FC-122-1001) and the exome prep was performed using KAPA Library Preparation Kit (Roche, #KK8201) using Agilent’s SureSelect Human All Exon V5 backbone with custom probes. The targeted DNA panel developed by Ashion Analytics (formerly known as the GEM Cancer panel) consisted of exonic probes against 541 cancer genes. Both panel and WXS assays contained 44,000 probes across evenly spaced genomic loci used for genome-wide copy number analysis. For the panel, additional probes tiled across intronic regions of 22 known tumor suppressor genes and 22 genes involved in common cancer translocations for structural analysis. All extractions and library preparations were performed according to manufacturer’s instructions.

#### CBTN samples

Blood, tissue, and cell line DNA/RNA extractions were performed at the Biorepository Core at CHOP. Briefly, 10-20 mg frozen tissue, 0.4-1ml of blood, or 2e6 cells pellet was used for extractions. Tissues were lysed using a Qiagen TissueLyser II (Qiagen) with 2×30 sec at 18Hz settings using 5 mm steel beads (cat# 69989, Qiagen). Both tissue and cell pellets processes included a CHCl3 extraction and were run on the QIACube automated platform (Qiagen) using the AllPrep DNA/RNA/miRNA Universal kit (cat# 80224, Qiagen). Blood was thawed and treated with RNase A (cat#, 19101, Qiagen); 0.4-1ml was processed using the Qiagen QIAsymphony automated platform (Qiagen) using the QIAsymphony DSP DNA Midi Kit (cat# 937255, Qiagen). DNA and RNA quantity and quality was assessed by PerkinElmer DropletQuant UV-VIS spectrophotometer (PerkinElmer) and an Agilent 4200 TapeStation (Agilent, USA) for RIN and DIN (RNA Integrity Number and DNA Integrity Number, respectively). The NantHealth Sequencing Center, BGI at CHOP, or the Genomic Clinical Core at Sidra Medical and Research Center performed library preparation and sequencing. BGI at CHOP and Sidra Medical and Research Center used in house, center-specific workflows for sample preparation. At NantHealth Sequencing Center, DNA sequencing libraries were prepared for tumor and matched-normal DNA using the KAPA HyperPrep kit (cat# 08098107702, Roche), and tumor RNA-Seq libraries were prepared using KAPA Stranded RNA-Seq with RiboErase kit (cat# 07962304001, Roche).

#### Data generation

NantHealth and Sidra performed 2×150 bp WGS on paired tumor (∼60X) and constitutive DNA (∼30X) samples on an Illumina X/400. BGI at CHOP performed 2×100 bp WGS sequenced at 60X depth for both tumor and normal samples. NantHealth performed ribosomal-depleted whole transcriptome stranded RNA-Seq to an average depth of 200M. BGI at CHOP performed poly-A or ribosomal-depleted whole transcriptome stranded RNA-Seq to an average depth of 100M. The Translational Genomic Research Institute (TGEN; Phoenix, AZ) performed paired tumor (∼200X) and constitutive whole exome sequencing (WXS) or targeted DNA panel (panel) and poly-A selected RNA-Seq (∼200M reads) for PNOC tumor samples. The panel tumor sample was sequenced to 470X, and the normal panel sample was sequenced to 308X. PNOC 2×100 bp WXS and RNA-Seq libraries were sequenced on an Illumina HiSeq 2500.

#### DNA WGS Alignment

We used `BWA-MEM`^74^ to align paired-end DNA-seq reads to the version 38 patch release 12 of the *Homo sapiens* genome reference, obtained as a FASTA file from UCSC (see **Key Resources Table**). Next, we used the Broad Institute’s Best Practices^75^ to process Binary Alignment/Map files (BAMs) in preparation for variant discovery. We marked duplicates using `SAMBLASTER`^76^, and we merged and sorted BAMs using `Sambamba`^77^ We used the `BaseRecalibrator` submodule of the Broad’s Genome Analysis Tool Kit `GATK`^78^ to process BAM files. Lastly, for normal/germline input, we used the `GATK HaplotypeCaller`^79^ submodule on the recalibrated BAM to generate a genomic variant call format (GVCF) file. This file is used as the basis for germline calling, described in the **SNV calling for B-allele Frequency (BAF) generation** section.

We obtained references from the <u>Broad Genome References on AWS</u> bucket with a general description of references at https://s3.amazonaws.com/broad-references/broad-references-readme.html.

#### Quality Control of Sequencing Data

To confirm sample matches and remove mis-matched samples from the dataset, we performed `NGSCheckMate`^80^ on matched tumor/normal CRAM files. Briefly, we processed CRAMs using `BCFtools` to filter and call 20k common single nucleotide polymorphisms (SNPs) using default parameters. We used the resulting VCFs to run `NGSCheckMate`. Per `NGSCheckMate` author recommendations, we used <= 0.61 as a correlation coefficient cutoff at sequencing depths > 10 to predict mis-matched samples. We determined RNA-Seq read strandedness by running the `infer_experiment.py` script from `RNA-SeQC`^81^ on the first 200k mapped reads. We removed any samples whose calculated strandedness did not match strandedness information provided by the sequencing center. We required that at least 60% of RNA-Seq reads mapped to the human reference for samples to be included in analysis.

### Germline Variant Calling

#### SNP calling for B-allele Frequency (BAF) generation

We performed germline haplotype calls using the `GATK` Joint Genotyping Workflow on individual GVCFs from the normal sample alignment workflow. Using only SNPs, we applied the `GATK` generic hard filter suggestions to the VCF, with an additional requirement of 10 reads minimum depth per SNP. We used the filtered VCF as input to `Control-FREEC` and `CNVkit` (below) to generate B-allele frequency (BAF) files. This single-sample workflow is available in the <u>D3b</u> <u>GitHub repository</u>. References can be obtained from the <u>Broad Genome References on AWS</u> bucket, and a general description of references can be found at https://s3.amazonaws.com/broad-references/broad-references-readme.html.

#### Assessment of germline variant pathogenicity

For patients with hypermutant samples, we first added population frequency of germline variants using `ANNOVAR`^82^ and pathogenicity scoring from ClinVar^83^ using `SnpSift`^84^. We then filtered for variants with read depth >= 15, variant allele fraction >= 0.20, and which were observed at < 0.1% allele frequency across each population in the Genome Aggregation Database (see **Key Resources Table**). Finally, we retained variants in genes included in the KEGG MMR gene set (see **Key Resources Table**), *POLE*, and/or *TP53* which were ClinVar-annotated as pathogenic (P) or likely pathogenic (LP) with review status of >= 2 stars. All P/LP variants were manually reviewed by an interdisciplinary team of scientists, clinicians, and genetic counselors. This workflow is available in the <u>D3b GitHub repository</u>.

### Somatic Mutation Calling

#### SNV and indel calling

For PBTA samples, we used four variant callers to call SNVs and indels from panel, WXS, and WGS data: `Strelka2`^85^, `Mutect2`^86^, `Lancet`^87^, and `VarDictJava`^88^. `VarDictJava`-only calls were not retained since ∼ 39M calls with low VAF were uniquely called and may be potential false positives. (∼1.2M calls were called by `Mutect2`, `Strelka2`, and `Lancet` and included consensus CNV calling as described below.) We used only `Strelka2`, `Mutect2` and `Lancet` to analyze WXS samples from TCGA. TCGA samples were captured using various WXS target capture kits and we downloaded the BED files from the `GDC portal`. The manufacturers provided the input interval BED files for both panel and WXS data for PBTA samples. We padded all panel and WXS BED files were by 100 bp on each side for `Strelka2`, `Mutect2`, and `VarDictJava` runs and by 400 bp for the `Lancet` run. For WGS calling, we utilized the non-padded BROAD Institute interval calling list `wgs_calling_regions.hg38.interval_list`, comprised of the full genome minus N bases, unless otherwise noted below. We ran `Strelka2`^85^ using default parameters for canonical chromosomes (chr1-22, X,Y,M), as recommended by the authors, and we filtered the final `Strelka2` VCF for PASS variants. We ran `Mutect2` from `GATK` according to Broad best practices outlined from their Workflow Description Language (WDL), and we filtered the final `Mutect2` VCF for PASS variants. To manage memory issues, we ran `VarDictJava`^88^ using 20 Kb interval chunks of the input BED, padded by 100 bp on each side, such that if an indel occurred in between intervals, it would be captured. Parameters and filtering followed <u>BCBIO</u> <u>standards</u> except that variants with a variant allele frequency (VAF) >= 0.05 (instead of >= 0.10) were retained. The 0.05 VAF increased the true positive rate for indels and decreased the false positive rate for SNVs when using `VarDictJava` in consensus calling. We filtered the final `VarDictJava` VCF for PASS variants with `TYPE=StronglySomatic`. We ran `Lancet` using default parameters, except for those noted below. For input intervals to `Lancet` WGS, we created a reference BED from only the UTR, exome, and start/stop codon features of the GENCODE 31 reference, augmented as recommended with PASS variant calls from `Strelka2` and `Mutect2`. We then padded these intervals by 300 bp on each side during `Lancet` variant calling. Per recommendations for WGS samples, we augmented the Lancet input intervals described above with PASS variant calls from `Strelka2` and `Mutect2` as validation^89^.

#### VCF annotation and MAF creation

We normalized INDELs with `bcftools norm` on all PASS VCFs using the `kfdrc_annot_vcf_sub_wf.cwl` subworkflow, release v3 (See Table S5). The Ensembl Variant Effect Predictor (`VEP`)^90^, reference release 93, was used to annotate variants and bcftools was used to add population allele frequency (AF) from gnomAD^91^. We annotated SNV and INDEL hotspots from v2 of Memorial Sloan Kettering Cancer Center’s (MSKCC) database (See **Key Resources Table**) as well as the *TERT* promoter mutations C228T and C250T^92^. We annotated SNVs by matching amino acid position (`Protein_position` column in MAF file) with SNVs in the MSKCC database, we matched splice sites to `HGVSp_Short` values in the MSKCC database, and we matched INDELs based on amino acid present within the range of INDEL hotspots values in the MSKCC database. We removed non-hotspot annotated variants with a normal depth less than or equal to 7 and/or gnomAD allele frequency (AF) greater than 0.001 as potential germline variants. We matched *TERT* promoter mutations using hg38 coordinates as indicated in ref.^92^: C228T occurs at 5:1295113 is annotated as existing variant `s1242535815`, `COSM1716563`, or `COSM1716558`, and is 66 bp away from the TSS; C250T occurs at Chr5:1295135, is annotated as existing variant `COSM1716559`, and is 88 bp away from the TSS. We retained variants annotated as `PASS` or `HotSpotAllele=1` in the final set, and we created MAFs using MSKCC’s `vcf2maf` tool.

#### Gather SNV and INDEL Hotspots

We retained all variant calls from `Strelka2`, `Mutect2`, or `Lancet` that overlapped with an SNV or INDEL hotspot in a hotspot-specific MAF file, which we then used for select analyses as described below.

#### Consensus SNV Calling

Our SNV calling process led to separate sets of predicted mutations for each caller. We considered mutations to describe the same change if they were identical for the following MAF fields: `Chromosome`, `Start_Position`, `Reference_Allele`, `Allele`, and `Tumor_Sample_Barcode`. `Strelka2` does not call multinucleotide variants (MNV), but instead calls each component SNV as a separate mutation, so we separated MNV calls from `Mutect2` and `Lancet` into consecutive SNVs before comparing them to `Strelka2` calls. We examined VAFs produced by each caller and compared their overlap with each other (Figure S2). `VarDictJava` calls included many variants that were not identified by other callers (Figure S2C), while the other callers produced results that were relatively consistent with one another. Many of these `VarDictJava`-specific calls were variants with low allele frequency (Figure S2B). We therefore derived consensus mutation calls as those shared among the other three callers (`Strelka2`, `Mutect2`, and `Lancet`), and we did not further consider `VarDictJava` calls due to concerns it called a large number of false positives. This decision had minimal impact on results because `VarDictJava` also identified nearly every mutation that the other three callers identified, in addition to many unique mutations.

#### Somatic Copy Number Variant Calling (WGS samples only)

We used `Control-FREEC`^93, 94^ and `CNVkit`^95^ for copy number variant calls. For both algorithms, the `germline_sex_estimate` (described below) was used as input for sample sex and germline variant calls (above) were used as input for BAF estimation. `Control-FREEC` was run on human genome reference hg38 using the optional parameters of a 0.05 coefficient of variation, ploidy choice of 2-4, and BAF adjustment for tumor-normal pairs. `Theta2`^96^ used `VarDictJava` germline and somatic calls, filtered on PASS and strongly somatic, to infer tumor purity. `Theta2` purity was added as an optional parameter to `CNVkit` to adjust copy number calls. `CNVkit` was run on human genome reference hg38 using the optional parameters of Theta2 purity and BAF adjustment for tumor-normal pairs. We used `GISTIC`^97^ on the `CNVkit` and the consensus CNV segmentation files to generate gene-level copy number abundance (Log R Ratio) as well as chromosomal arm copy number alterations using the parameters specified in the (`run-gistic` analysis module in the OpenPBTA Analysis repository).

#### Consensus CNV Calling

For each caller and sample, we called CNVs based on consensus among `Control-FREEC`^93, 94^, `CNVkit`^95^, and `Manta`^98^. We specifically included CNVs called significant by `Control-FREEC` (p-value < 0.01) and `Manta` calls that passed all filters in consensus calling. We removed sample and consensus caller files with more than 2,500 CNVs because we expected these to be noisy and derive poor quality samples based on cutoffs used in `GISTIC`^97^. For each sample, we included the regions in the final consensus set: 1) regions with reciprocal overlap of 50% or more between at least two of the callers; 2) smaller CNV regions in which more than 90% of regions are covered by another caller. We did not include any copy number alteration called by a single algorithm in the consensus file. We defined copy number as `NA` for any regions that had a neutral call for the samples included in the consensus file. We merged CNV regions within 10,000 bp of each other with the same direction of gain or loss into single region. We filtered out any CNVs that overlapped 50% or more with immunoglobulin, telomeric, centromeric, segment duplicated regions, or that were shorter than 3000 bp.

#### Somatic Structural Variant Calling (WGS samples only)

We used `Manta`^98^ for structural variant (SV) calls, and we limited to regions used in `Strelka2`. The hg38 reference for SV calling used was limited to canonical chromosome regions. We used `AnnotSV`^99^ to annotate `Manta` output. All associated workflows are available in the <u>workflows</u> <u>GitHub repository</u>.

### Gene Expression

#### Abundance Estimation

We used `STAR`^100^ to align paired-end RNA-seq reads, and we used the associated alignment for all subsequent RNA analysis. We used Ensembl GENCODE 27 “Comprehensive gene annotation” (see **Key Resources Table**) as a reference. We used `RSEM`^101^ for both FPKM and TPM transcript- and gene-level quantification.

#### Gene Expression Matrices with Unique HUGO Symbols

To enable downstream analyses, we next identified gene symbols that map to multiple Ensembl gene identifiers (in GENCODE v27, 212 gene symbols map to 1866 Ensembl gene identifiers), known as multi-mapped gene symbols, and ensured unique mappings (`collapse-rnaseq` analysis module in the OpenPBTA Analysis repository). To this end, we first removed genes with no expression from the `RSEM` abundance data by requiring an FPKM > 0 in at least 1 sample across the PBTA cohort. We computed the mean FPKM across all samples per gene.

For each multi-mapped gene symbol, we chose the Ensembl identifier corresponding to the maximum mean FPKM, using the assumption that the gene identifier with the highest expression best represented the expression of the gene. After collapsing gene identifiers, 46,400 uniquely-expressed genes remained in the poly-A dataset, and 53,011 uniquely-expressed genes remained in the stranded dataset.

#### Gene fusion detection

We set up `Arriba`^102^ and `STAR-Fusion`^103^ fusion detection tools using CWL on CAVATICA. For both of these tools, we used aligned BAM and chimeric SAM files from `STAR` as inputs and `GRCh38_gencode_v27` GTF for gene annotation. We ran `STAR-Fusion` with default parameters and annotated all fusion calls with the `GRCh38_v27_CTAT_lib_Feb092018.plug-n-play.tar.gz` file from the `STAR-Fusion` release. For `Arriba`, we used a blacklist file `blacklist_hg38_GRCh38_2018-11-04.tsv.gz` from the `Arriba` release to remove recurrent fusion artifacts and transcripts present in healthy tissue. We provided `Arriba` with strandedness information for stranded samples, or we set it to auto-detection for poly-A samples. We used `FusionAnnotator` on `Arriba` fusion calls to harmonize annotations with those of `STAR-Fusion`. The RNA expression and fusion workflows can be found in the D3b GitHub repository. The `FusionAnnotator` workflow we used for this analysis can be found in the D3b GitHub <u>repository</u>.

### QUANTIFICATION AND STATISTICAL ANALYSIS

#### Recurrently mutated genes and co-occurrence of gene mutations (interaction-plots analysis module)

Using the consensus SNV calls, we identified genes that were recurrently mutated in the OpenPBTA cohort, including nonsynonymous mutations with a VAF > 5% among the set of independent samples. We used `VEP`^90^ annotations, including “High” and “Moderate” consequence types as defined in the R package `Maftools`^104^, to determine the set of nonsynonymous mutations. For each gene, we then tallied the number of samples that had at least one nonsynonymous mutation.

For genes that contained nonsynonymous mutations in multiple samples, we calculated pairwise mutation co-occurrence scores. This score was defined as *I*(–log_10_(*P*)) where *I* is 1 when the odds ratio is > 1 (indicating co-occurrence), and −1 when the odds ratio is < 1 (indicating mutual exclusivity), with *P* defined by Fisher’s Exact Test.

#### Focal Copy Number Calling (focal-cn-file-preparation analysis module)

We added the ploidy inferred via `Control-FREEC` to the consensus CNV segmentation file and used the ploidy and copy number values to define gain and loss values broadly at the chromosome level. We used `bedtools coverage`^105^ to add cytoband status using the UCSC cytoband file^106^ (See **Key Resources Table**). The output status call fractions, which are values of the loss, gain, and callable fractions of each cytoband region, were used to define dominant status at the cytoband-level. We calculated the weighted means of each status call fraction using band length. We used the weighted means to define the dominant status at the chromosome arm-level.

A status was considered dominant if more than half of the region was callable and the status call fraction was greater than 0.9 for that region. We adopted this 0.9 threshold to ensure that the dominant status fraction call was greater than the remaining status fraction calls in a region.

We aimed to define focal copy number units to avoid calling adjacent genes in the same cytoband or arm as copy number losses or gains where it would be more appropriate to call the broader region a loss or gain. To determine the most focal units, we first considered the dominant status calls at the chromosome arm-level. If the chromosome arm dominant status was callable but not clearly defined as a gain or loss, we instead included the cytoband-level status call. Similarly, if a cytoband dominant status call was callable but not clearly defined as a gain or loss, we instead included gene-level status call. To obtain the gene-level data, we used the `IRanges` package in R^107^ to find overlaps between the segments in the consensus CNV file and the exons in the GENCODE v27 annotation file (See **Key Resources Table**). If the copy number value was 0, we set the status to “deep deletion”. For autosomes only, we set the status to “amplification” when the copy number value was greater than two times the ploidy value. We plotted genome-wide gains and losses in (Figure S3B) using the R package `ComplexHeatmap`^108^.

#### Breakpoint Density (WGS samples only; chromosomal-instability analysis module)

We defined breakpoint density as the number of breaks per genome or exome per sample. For Manta SV calls, we filtered to retain “PASS” variants and used breakpoints from the algorithm. For consensus CNV calls, if |log2 ratio| > log2(1), we annotated the segment as a break. We then calculated breakpoint density as:

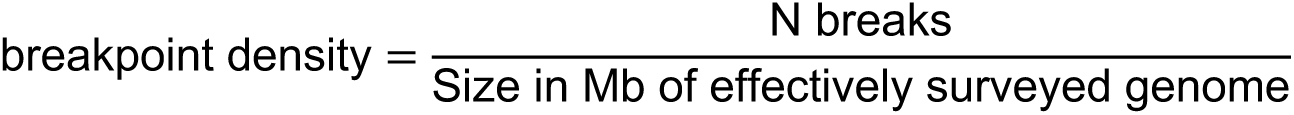

#### Chromothripsis Analysis (WGS samples only; chromothripsis analysis module)

Considering only chromosomes 1-22 and X, we identified candidate chromothripsis regions in the set of independent tumor WGS samples with ShatterSeek^109^, using Manta SV calls that passed all filters and consensus CNV calls. We modified the consensus CNV data to fit `ShatterSeek` input requirements as follows: we set CNV-neutral or excluded regions as the respective sample’s ploidy value from `Control-FREEC`, and we then merged consecutive segments with the same copy number value. We classified candidate chromothripsis regions as high- or low-confidence using the statistical criteria described by the `ShatterSeek` authors.

#### Immune Profiling and Deconvolution (immune-deconv analysis module)

We used the R package `immunedeconv`^110^ with the method `quanTIseq`^111^ to deconvolute various immune cell types across tumors from the PBTA cohort in the stranded and poly-A collapsed FPKM RNA-seq datasets (`immune-deconv` <u>analysis module</u>). The `quanTIseq` deconvolution method directly estimates absolute fractions of 10 immune cell types that represent inferred proportions of the cell types in the mixture. Therefore, we utilized `quanTIseq` for inter-sample, intra-sample, and inter-histology score comparisons.

#### Gene Set Variation Analysis (gene-set-enrichment-analysis analysis module)

We performed Gene Set Variation Analysis (GSVA) on collapsed, log2-transformed RSEM FPKM data using the `GSVA` Bioconductor package^112^. We specified the parameter `mx.diff=TRUE` to obtain Gaussian-distributed scores for each of the MSigDB hallmark gene sets^113^. We compared GSVA scores among histology groups using ANOVA and subsequent Tukey tests; p-values were Bonferroni-corrected for multiple hypothesis testing. We plotted scores by cancer group using the `ComplexHeatmap` R package (Figure 5B)^108^.

#### Transcriptomic Dimension Reduction (transcriptomic-dimension-reduction analysis module)

We applied Uniform Manifold Approximation and Projection (UMAP)^114^ to log2-transformed FPKM data using the `umap` R package (See **Key Resources Table**). We set the number of neighbors to 15.

#### Fusion prioritization (fusion_filtering analysis module)

We performed artifact filtering and additional annotation on fusion calls to prioritize putative oncogenic fusions. Briefly, we considered all in-frame and frameshift fusion calls with at least one junction read and at least one gene partner expressed (TPM > 1) to be true calls. If a fusion call had a large number of spanning fragment reads compared to junction reads (spanning fragment minus junction read greater than ten), we removed these calls as potential false positives. We prioritized a union of fusion calls as true calls if the fused genes were detected by both callers, the same fusion was recurrent within a broad histology grouping (> 2 samples), or the fusion was specific to the given broad histology. If either 5’ or 3’ genes fused to more than five different genes within a sample, we removed these calls as potential false positives. We annotated putative driver fusions and prioritized fusions based on partners containing known <u>kinases</u>, <u>oncogenes</u>, <u>tumor suppressors</u>, curated transcription factors^115^, <u>COSMIC genes</u>, and/or known <u>TCGA fusions</u> from curated references. Based on pediatric cancer literature review, we added *MYBL1*^116^, *SNCAIP*^117^, *FOXR2*^118^, *TTYH1*^119^, and *TERT*^120–123^ to the oncogene list, and we added *BCOR*^118^ and *QKI*^124^ to the tumor suppressor gene list.

#### Oncoprint figure generation (oncoprint-landscape analysis module)

We used `Maftools`^104^ to generate oncoprints depicting the frequencies of canonical somatic gene mutations, CNVs, and fusions for the top 20 genes mutated across primary tumors within broad histologies of the OpenPBTA dataset. We collated canonical genes from the literature for low-grade astrocytic tumors^25^, embryonal tumors^26, 28, 29, 125, 126^, diffuse astrocytic and oligodendroglial tumors^15, 22, 30, 31^, and other tumors: ependymal tumors, craniopharyngiomas, neuronal-glial mixed tumors, histiocytic tumors, chordoma, meningioma, and choroid plexus tumors^127–136^.

#### Mutational Signatures (mutational-signatures analysis module)

We obtained weights (i.e., exposures) for signature sets using the `deconstructSigs` R package function `whichSignatures()`^137^ from consensus SNVs with the BSgenome.Hsapiens.UCSC.hg38 annotations (see **Key Resources Table**). Specifically, we estimated signature weights across samples for eight signatures previously identified in the Signal reference set of signatures (“RefSig”) as associated with adult central nervous system (CNS) tumors^36^. These eight RefSig signatures are 1, 3, 8, 11, 18, 19, N6, and MMR2. Weights for signatures fall in the range zero to one inclusive. `deconstructSigs` estimates the weights for each signature across samples and allows for a proportion of unassigned weights referred to as “Other” in the text. These results do not include signatures with small contributions; `deconstructSigs` drops signature weights that are less than 6%^137^. We plotted mutational signatures for patients with hypermutant tumors (Figure 4E) using the R package `ComplexHeatmap`^108^.

#### Tumor Mutation Burden (snv-callers analysis module)

We consider tumor mutation burden (TMB) to be the number of consensus SNVs per effectively surveyed base of the genome. We considered base pairs to be effectively surveyed if they were in the intersection of the genomic ranges considered by the callers used to generate the consensus and where appropriate, regions of interest, such as coding sequences. We calculated TMB as:

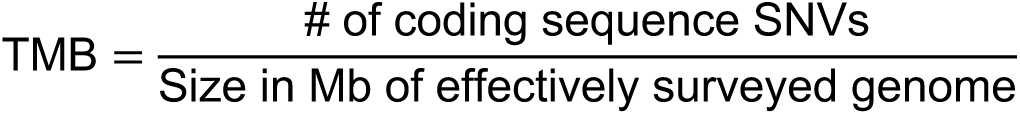

We used the total number coding sequence consensus SNVs for the numerator and the size of the intersection of the regions considered by `Strelka2` and `Mutect2` with coding regions (CDS from GENCODE v27 annotation, see **Key Resources Table**) as the denominator.

### Clinical Data Harmonization

#### WHO Classification of Disease Types

Table S1 contains a README, along with sample technical, clinical, and additional metadata used for this study.

#### Molecular Subtyping

We performed molecular subtyping on tumors in the OpenPBTA to the extent possible. The `molecular_subtype` field in `pbta-histologies.tsv` contains molecular subtypes for tumor types selected from `pathology_diagnosis` and `pathology_free_text_diagnosis` fields as described below, following World Health Organization 2016 classification criteria^21^.

Medulloblastoma (MB) subtypes SHH, WNT, Group 3, and Group 4 were predicted using the consensus of two RNA expression classifiers: `MedulloClassifier`^61^ and `MM2S`^60^ on the RSEM FPKM data (`molecular-subtyping-MB` analysis module).

High-grade glioma (HGG) subtypes were derived (`molecular-subtyping-HGG` analysis module) using the following criteria:

1. If any sample contained an *H3F3A* p.K28M, *HIST1H3B* p.K28M, *HIST1H3C* p.K28M, or *HIST2H3C* p.K28M mutation and no *BRAF* p.V600E mutation, it was subtyped as `DMG, H3 K28`.
2. If any sample contained an *HIST1H3B* p.K28M, *HIST1H3C* p.K28M, or *HIST2H3C* p.K28M mutation and a *BRAF* p.V600E mutation, it was subtyped as `DMG, H3 K28, BRAF V600E`.
3. If any sample contained an *H3F3A* p.G35V or p.G35R mutation, it was subtyped as`HGG, H3 G35`.
4. If any high-grade glioma sample contained an *IDH1* p.R132 mutation, it was subtyped as `HGG, IDH`.
5. If a sample was initially classified as HGG, had no defining histone mutations, and a BRAF p.V600E mutation, it was subtyped as `BRAF V600E`.
6. All other high-grade glioma samples that did not meet any of these criteria were subtyped as `HGG, H3 wildtype`.

Embryonal tumors were included in non-MB and non-ATRT embryonal tumor subtyping (`molecular-subtyping-embryonal` analysis module) if they met any of the following criteria:

1. A *TTYH1* (5’ partner) fusion was detected.
2. A *MN1* (5’ partner) fusion was detected, with the exception of `MN1::PATZ1` since it is an entity separate of CNS HGNET-MN1 tumors^138^.
3. Pathology diagnoses included “Supratentorial or Spinal Cord PNET” or “Embryonal Tumor with Multilayered Rosettes”.
4. A pathology diagnosis of “Neuroblastoma”, where the tumor was not indicated to be peripheral or metastatic and was located in the CNS.
5. Any sample with “embryonal tumor with multilayer rosettes, ros (who grade iv)”, “embryonal tumor, nos, congenital type”, “ependymoblastoma” or “medulloepithelioma” in pathology free text.

Non-MB and non-ATRT embryonal tumors identified with the above criteria were further subtyped (`molecular-subtyping-embryonal` analysis module) using the criteria below^139–142^.

1. Any RNA-seq biospecimen with *LIN28A* overexpression, plus a *TYH1* fusion (5’ partner) with a gene adjacent or within the C19MC miRNA cluster and/or copy number amplification of the C19MC region was subtyped as `ETMR, C19MC-altered` (Embryonal tumor with multilayer rosettes, chromosome 19 miRNA cluster altered)^119, 143^.
2. Any RNA-seq biospecimen with *LIN28A* overexpression, a *TTYH1* fusion (5’ partner) with a gene adjacent or within the C19MC miRNA cluster but no evidence of copy number amplification of the C19MC region was subtyped as `ETMR, NOS` (Embryonal tumor with multilayer rosettes, not otherwise specified)^119, 143^.
3. Any RNA-seq biospecimen with a fusion having a 5’ *MN1* and 3’ *BEND2* or *CXXC5* partner were subtyped as `CNS HGNET-MN1` [Central nervous system (CNS) high-grade neuroepithelial tumor with *MN1* alteration].
4. Non-MB and non-ATRT embryonal tumors with internal tandem duplication (as defined in^144^) of *BCOR* were subtyped as `CNS HGNET-BCOR` (CNS high-grade neuroepithelial tumor with *BCOR* alteration).
5. Non-MB and non-ATRT embryonal tumors with over-expression and/or gene fusions in *FOXR2* were subtyped as `CNS NB-FOXR2` (CNS neuroblastoma with *FOXR2* activation).
6. Non-MB and non-ATRT embryonal tumors with *CIC::NUTM1* or other *CIC* fusions, were subtyped as `CNS EFT-CIC` (CNS Ewing sarcoma family tumor with *CIC* alteration)^118^
7. Non-MB and non-ATRT embryonal tumors that did not fit any of the above categories were subtyped as `CNS Embryonal, NOS` (CNS Embryonal tumor, not otherwise specified).

Neurocytoma subtypes central neurocytoma (CNC) and extraventricular neurocytoma (EVN) were assigned (`molecular-subtyping-neurocytoma` analysis module) based on the primary site of the tumor^145^. If the tumor’s primary site was “ventricles,” we assigned the subtype as CNC; otherwise, we assigned the subtype as EVN.

Craniopharyngiomas (CRANIO) were subtyped (`molecular-subtyping-CRANIO` analysis module) into adamantinomatous (`CRANIO, ADAM`), papillary (`CRANIO, PAP`) or undetermined (`CRANIO, To be classified`) based on the following criteria^146, 147^:

1. Craniopharyngiomas from patients over 40 years old with a *BRAF* p.V600E mutation were subtyped as `CRANIO, PAP`.
2. Craniopharyngiomas from patients younger than 40 years old with mutations in exon 3 of *CTNNB1* were subtyped as `CRANIO, ADAM`.
3. Craniopharyngiomas that did not fall into the above two categories were subtyped as

`CRANIO, To be classified`.

A molecular subtype of `EWS` was assigned to any tumor with a *EWSR1* fusion or with a `pathology_diagnosis` of `Ewings Sarcoma` (`molecular-subtyping-EWS` analysis module).

Low-grade gliomas (LGG) or glialneuronal tumors (GNT) were subtyped (`molecular-subtyping-LGAT` analysis module). based on SNV, fusion and CNV status based on^23^, and as described below.

1. If a sample contained a *NF1* somatic mutation, either nonsense or missense, it was subtyped as `LGG, NF1-somatic`.
2. If a sample contained *NF1* germline mutation, as indicated by a patient having the neurofibromatosis cancer predisposition, it was subtyped as `LGG, NF1-germline`.
3. If a sample contained the *IDH* p.R132 mutation, it was subtyped as `LGG, IDH`.
4. If a sample contained a histone p.K28M mutation in either *H3F3A*, *H3F3B*, *HIST1H3B*, *HIST1H3C*, or *HIST2H3C*, or if it contained a p.G35R or p.G35V mutation in *H3F3A*, it was subtyped as `LGG, H3`.
5. If a sample contained *BRAF* p.V600E or any other non-canonical *BRAF* mutations in the kinase (PK_Tyr_Ser-Thr) domain PF07714 (see **Key Resources Table**), it was subtyped as `LGG, BRAF V600E`.
6. If a sample contained `KIAA1549::BRAF` fusion, it was subtyped as `LGG, KIAA1549::BRAF`.
7. If a sample contained SNV or indel in either *KRAS*, *NRAS*, *HRAS*, *MAP2K1*, *MAP2K2*, *MAP2K1*, *ARAF*, *RAF1*, or non-kinase domain of *BRAF*, or if it contained *RAF1* fusion, or *BRAF* fusion that was not `KIAA1549::BRAF`, it was subtyped as `LGG, other MAPK`.
8. If a sample contained SNV in either *MET*, *KIT* or *PDGFRA*, or if it contained fusion in *ALK*, *ROS1*, *NTRK1*, *NTRK2*, *NTRK3* or *PDGFRA*, it was subtyped as `LGG, RTK`.
9. If a sample contained *FGFR1* p.N546K, p.K656E, p.N577, or p. K687 hotspot mutations, or tyrosine kinase domain tandem duplication (See **Key Resources Table**), or *FGFR1* or *FGFR2* fusions, it was subtyped as `LGG, FGFR`.
10. If a sample contained *MYB* or *MYBL1* fusion, it was subtyped as `LGG, MYB/MYBL1`.
11. If a sample contained focal CDKN2A and/or CDKN2B deletion, it was subtyped as `LGG, CDKN2A/B`.

For LGG tumors that did not have any of the above molecular alterations, if both RNA and DNA samples were available, it was subtyped as `LGG, wildtype`. Otherwise, if either RNA or DNA sample was unavailable, it was subtyped as `LGG, To be classified`.

If pathology diagnosis was `Subependymal Giant Cell Astrocytoma (SEGA)`, the `LGG` portion of molecular subtype was recoded to `SEGA`.

Lastly, for all LGG- and GNT-subtyped samples, if the tumors were glialneuronal in origin, based on `pathology_free_text_diagnosis` entries of `desmoplastic infantile`,`desmoplastic infantile ganglioglioma`, `desmoplastic infantile astrocytoma` or `glioneuronal`, each was recoded as follows: If pathology diagnosis is `Low-grade glioma/astrocytoma (WHO grade I/II)` or `Ganglioglioma`, the `LGG` portion of the molecular subtype was recoded to `GNT`.

Ependymomas (EPN) were subtyped (`molecular-subtyping-EPN` analysis module) into `EPN, ST RELA`, `EPN, ST YAP1`, `EPN, PF A` and `EPN, PF B` based on evidence for these molecular subgroups as described in Pajtler et al.^128^. Briefly, fusion, CNV and gene expression data were used to subtype EPN as follows:

1. Any tumor with fusions containing `RELA` as fusion partner, e.g., `C11orf95::RELA`, `LTBP3::RELA`, was subtyped as `EPN, ST RELA`.
2. Any tumor with fusions containing `YAP1` as fusion partner, such as `C11orf95::YAP1`, `YAP1::MAMLD1` and `YAP1::FAM118B`, was subtyped as `EPN, ST YAP1`.
3. Any tumor with the following molecular characterization would be subtyped as `EPN, PF A`:
  - *CXorf67* expression z-score of over 3
  - *TKTL1* expression z-score of over 3 and 1q gain
4. Any tumor with the following molecular characterization would be subtyped as `EPN, PF B`:
  - *GPBP17* expression z-score of over 3 and loss of 6q or 6p
  - *IFT46* expression z-score of over 3 and loss of 6q or 6p

Any tumor with the above molecular characteristics would be exclusively subtyped to the designated group.

For all other remaining EPN tumors without above molecular characteristics, they would be subtyped to `EPN, ST RELA` and `EPN, ST YAP1` in a non-exclusive way (e.g., a tumor could have both `EPN, ST RELA` and `EPN, ST YAP1` subtypes) if any of the following alterations were present.

1. Any tumor with the following alterations was assigned `EPN, ST RELA`:
  - `PTEN::TAS2R1` fusion
  - chromosome 9 arm (9p or 9q) loss
  - *RELA* expression z-score of over 3
  - *L1CAM* expression z-score of over 3
2. Any tumor with the following alterations was assigned `EPN, ST YAP1`:
  - `C11orf95::MAML2` fusion
  - chromosome 11 short arm (11p) loss
  - chromosome 11 long arm (11q) gain
  - *ARL4D* expression z-score of over 3
  - *CLDN1* expression z-score of over 3

After all relevant tumor samples were subtyped by the above molecular subtyping modules, the results from these modules, along with other clinical information (such as pathology diagnosis free text), were compiled in the `molecular-subtyping-pathology` module and integrated into the OpenPBTA data in the `molecular-subtyping-integrate` module.

#### TP53 Alteration Annotation (tp53_nf1_score analysis module)

We annotated *TP53* altered HGG samples as either `TP53 lost` or `TP53 activated` and integrated this within the molecular subtype. To this end, we applied a *TP53* inactivation classifier originally trained on TCGA pan-cancer data^38^ to the matched RNA expression data for each sample. Along with the *TP53* classifier scores, we collectively used consensus SNV and CNV, SV, and reference databases that list *TP53* hotspot mutations^148, 149^ and functional domains^150^ to determine *TP53* alteration status for each sample. We adopted the following rules for calling either `TP53 lost` or `TP53 activated`:

1. If a sample had either of the two well-characterized *TP53* gain-of-function mutations, p.R273C or p.R248W^39^, we assigned `TP53 activated` status.
2. Samples were annotated as `TP53 lost` if they contained i) a *TP53* hotspot mutation as defined by IARC *TP53* database or the MSKCC cancer hotspots database^148, 149^ (see also, **Key Resources Table**), ii) two *TP53* alterations, including SNV, CNV or SV, indicative of probable bi-allelic alterations; iii) one *TP53* somatic alteration, including

SNV, CNV, or SV or a germline *TP53* mutation indicated by the diagnosis of Li-Fraumeni syndrome (LFS)^151^, or iv) one germline *TP53* mutation indicated by LFS and the *TP53* classifier score for matched RNA-Seq was greater than 0.5.

#### Prediction of participants’ genetic sex

Participant metadata included a reported gender. We used WGS germline data, in concert with the reported gender, to predict participant genetic sex so that we could identify sexually dimorphic outcomes. This analysis may also indicate samples that may have been contaminated. We used the `idxstats` utility from `SAMtools`^152^ to calculate read lengths, the number of mapped reads, and the corresponding chromosomal location for reads to the X and Y chromosomes. We used the fraction of total normalized X and Y chromosome reads that were attributed to the Y chromosome as a summary statistic. We manually reviewed this statistic in the context of reported gender and determined that a threshold of less than 0.2 clearly delineated female samples. We marked fractions greater than 0.4 as predicted males, and we marked samples with values in the inclusive range 0.2-0.4 as unknown. We performed this analysis through <u>CWL</u> on CAVATICA. We added resulting calls to the histologies file under the column header `germline_sex_estimate`.

#### Selection of independent samples (independent-samples analysis module)

Certain analyses required that we select only a single representative specimen for each individual. In these cases, we identified a single specimen by prioritizing primary tumors and those with whole-genome sequencing available. If this filtering still resulted in multiple specimens, we randomly selected a single specimen from the remaining set.

#### Quantification of Telomerase Activity using Gene Expression Data (telomerase-activity-prediction analysis module)

We predicted telomerase activity of tumor samples using the recently developed `EXTEND` method^41^. Briefly, `EXTEND` estimates telomerase activity based on the expression of a 13-gene signature. We derived this signature by comparing telomerase-positive tumors and tumors with activated alternative lengthening of telomeres pathway, a group presumably negative of telomerase activity.

#### Survival models (survival-analysis analysis module)

We calculated overall survival (OS) as days since initial diagnosis and performed several survival analyses on the OpenPBTA cohort using the `survival` <u>R package</u>. We performed survival analysis for patients by HGG subtype using the Kaplan-Meier estimator^153^ and a log-rank test (Mantel-Cox test)^154^ on the different HGG subtypes. Next, we used multivariate Cox (proportional hazards) regression analysis^155^ to model the following: a) `tp53 scores + telomerase scores + extent of tumor resection + LGG group + HGG group`, in which `tp53 scores` and `telomerase scores` are numeric, `extent of tumor resection` is categorical, and `LGG group` and `HGG group` are binary variables indicating whether the sample is in either broad histology grouping, b) `tp53 scores + telomerase scores + extent of tumor resection` for each `cancer_group` with an N>=3 deceased patients (DIPG, DMG, HGG, MB, and EPN), and c) `quantiseq cell type fractions + CD274 expression + extent of tumor resection` for each `cancer_group` with an N>=3 deceased patients (DIPG, DMG, HGG, MB, and EPN), in which `quantiseq cell type fractions` and `CD274 expression` are numeric.

## KEY RESOURCES TABLE

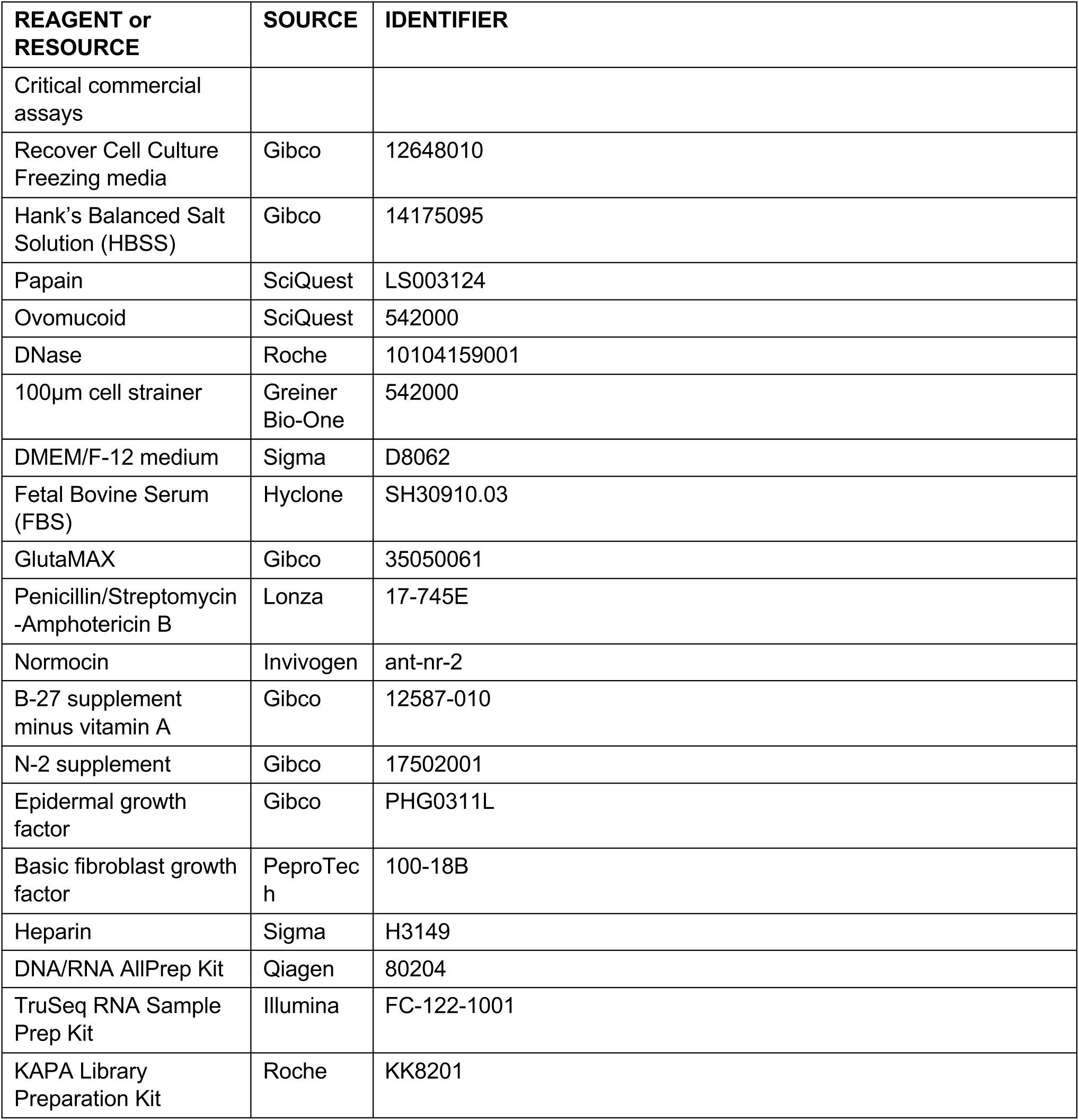
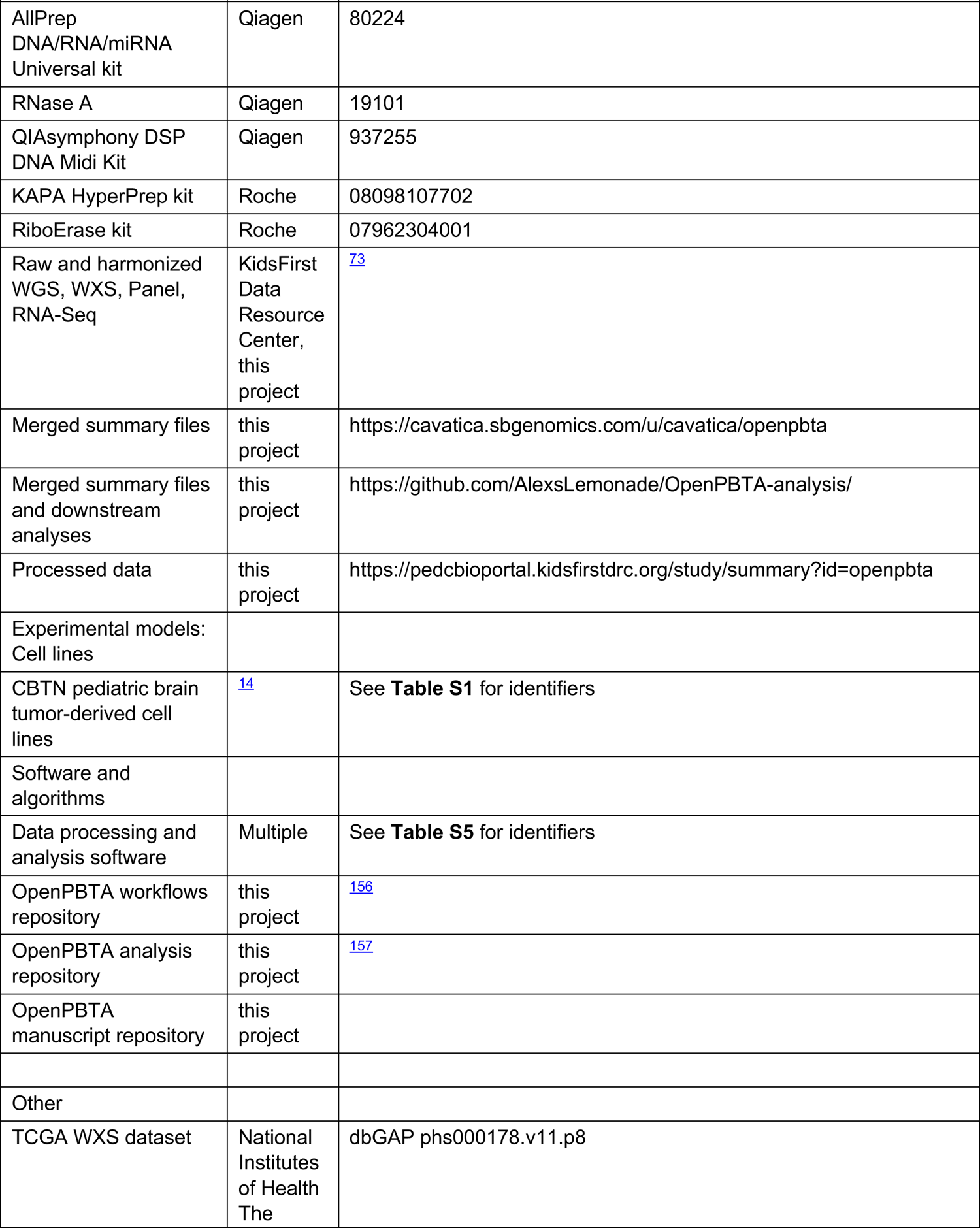
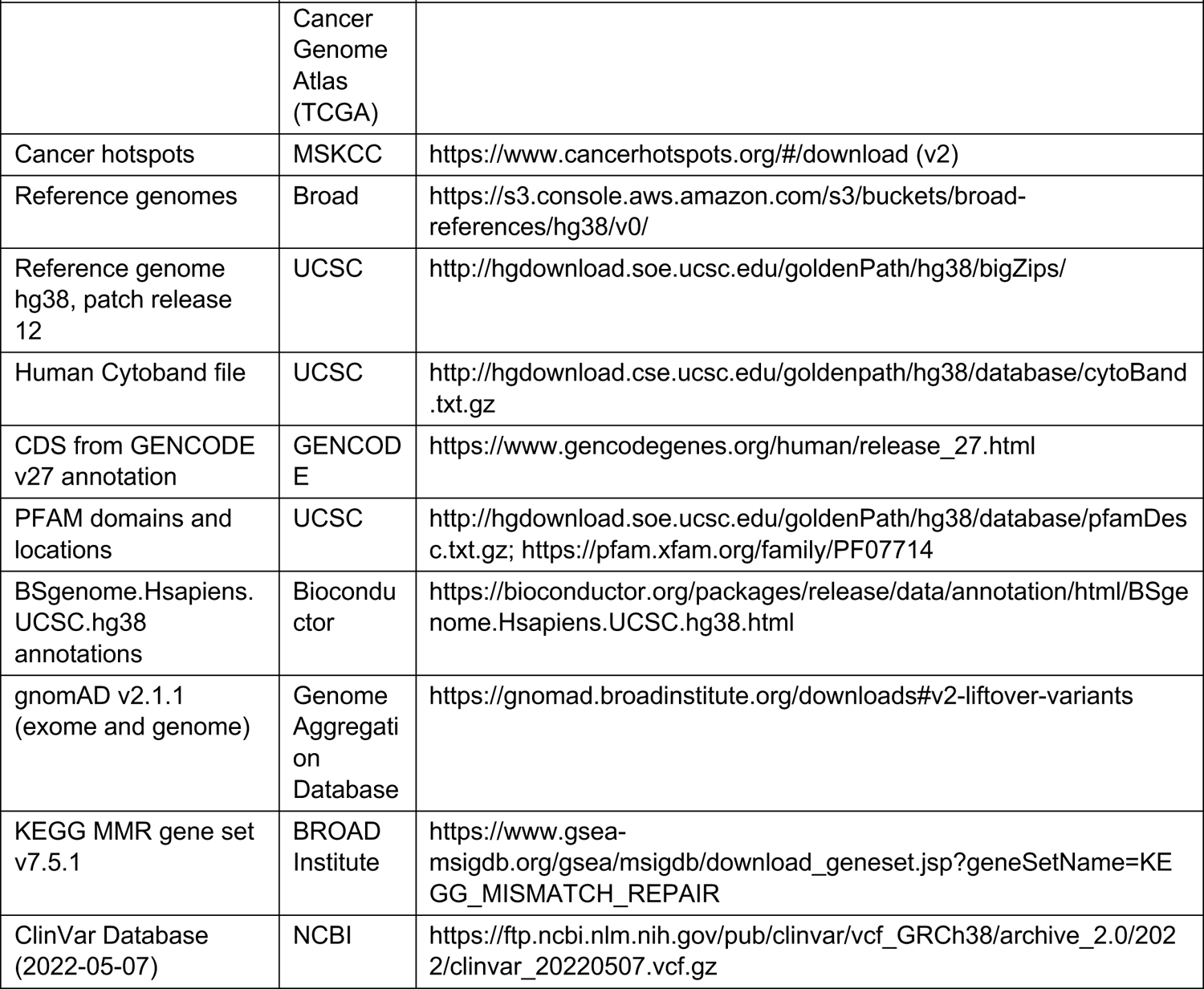

## Supplemental Information Titles and Legends

**Figure S1:**
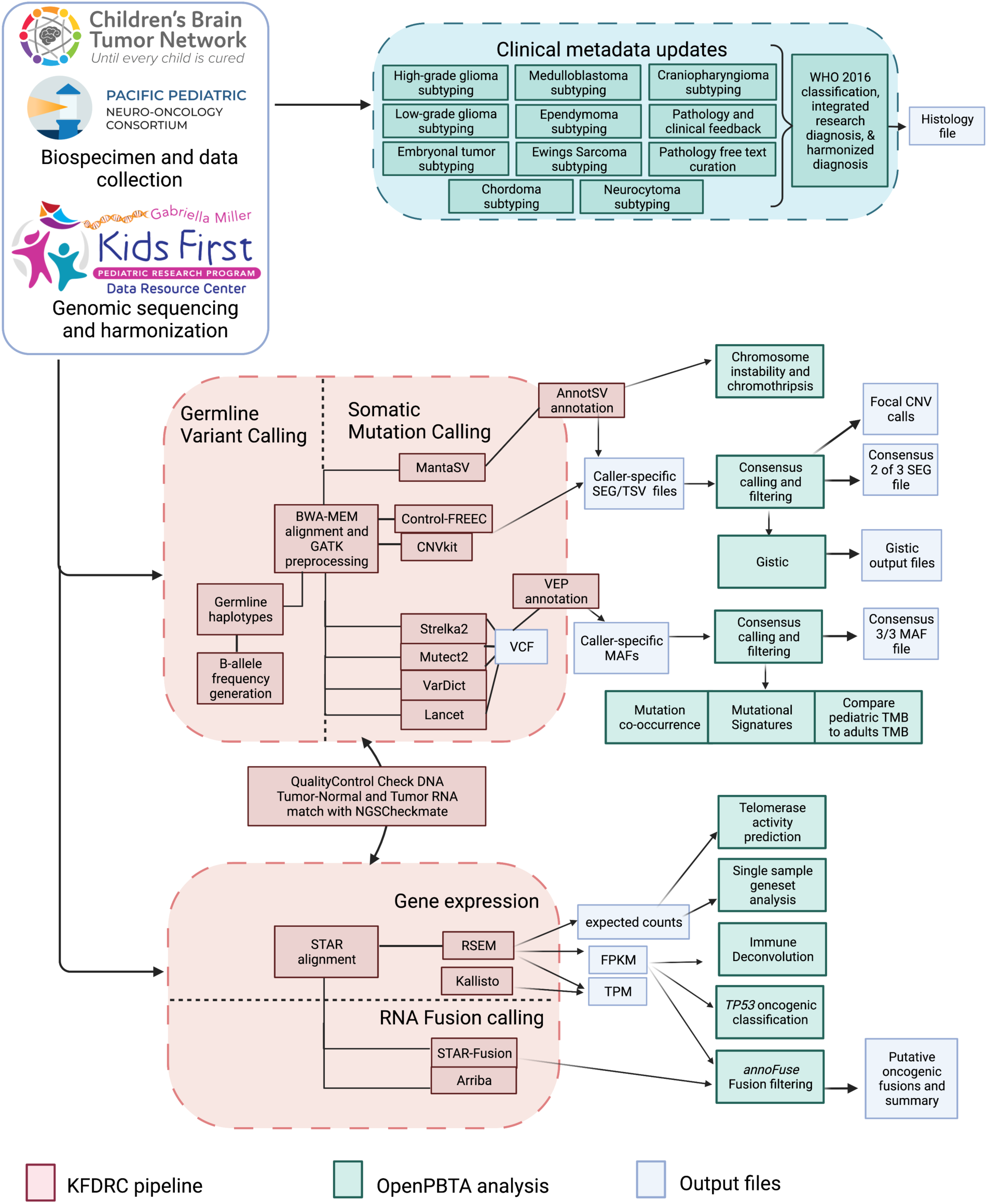
OpenPBTA Project Workflow, Related to Figure 1. Biospecimens and data were collected by CBTN and PNOC. Genomic sequencing and harmonization (orange boxes) were performed by the Kids First Data Resource Center (KFDRC). Analyses in the green boxes were performed by contributors of the OpenPBTA project. Output files are denoted in blue. Figure created with <u>BioRender.com</u>.

**Figure S2:**
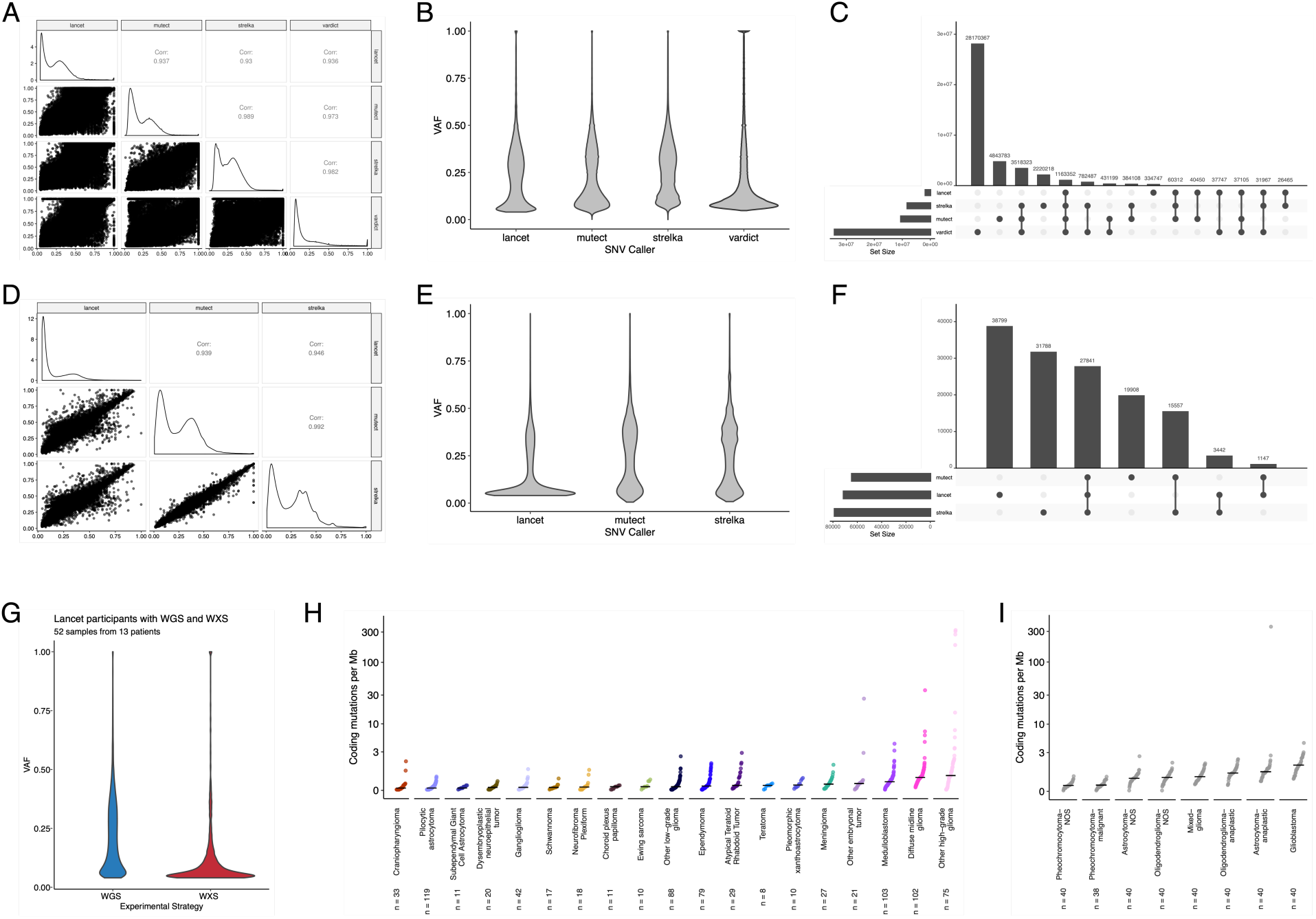
Validation of Consensus SNV calls and Tumor Mutation Burden, Related to Figures 2 and 3. Correlation (A) and violin (B) plots of mutation variant allele frequencies (VAFs) comparing the variant callers (Lancet, Strelka2, Mutect2, and VarDict) used for PBTA samples. Upset plot (C) showing overlap of variant calls. Correlation (D) and violin (E) plots of mutation variant allele frequencies (VAFs) comparing the variant callers (Lancet, Strelka2, and Mutect2) used for TCGA samples. Upset plot (F) showing overlap of variant calls. Violin plots (G) showing VAFs for Lancet calls performed on WGS and WXS from the same tumor (N = 52 samples from 13 patients). Cumulative distribution TMB plots for PBTA (H) and TCGA (I) tumors using consensus SNV calls.

**Figure S3:**
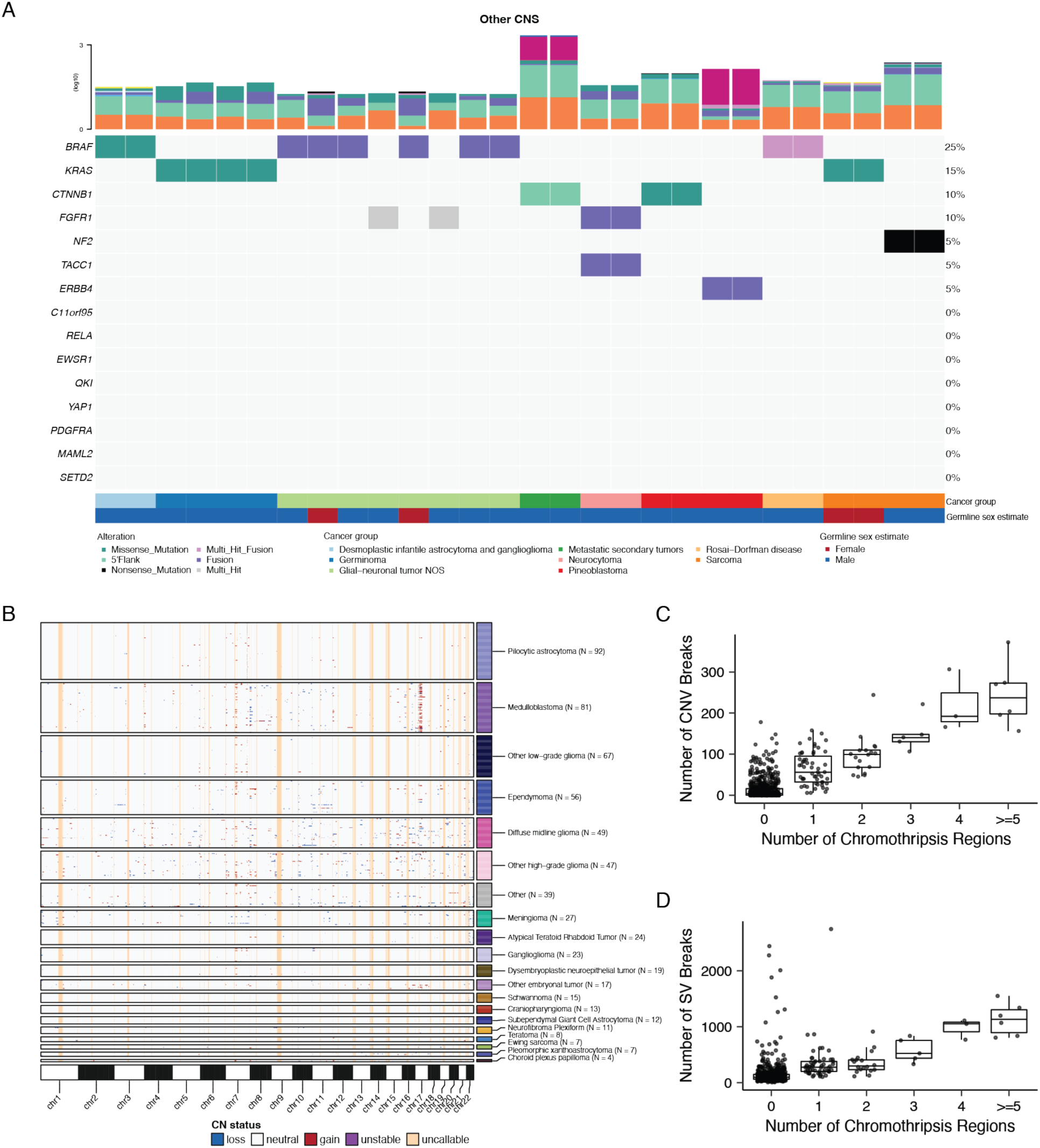
Genomic instability of pediatric brain tumors, Related to Figures 2 and 3. (A) Oncoprint of canonical somatic gene mutations, CNVs, fusions, and TMB (top bar plot) for the top 20 genes mutated across rare CNS tumors: desmoplastic infantile astrocytoma and ganglioglioma (N = 2), germinoma (N = 4), glial-neuronal NOS (N = 8), metastatic secondary tumors (N = 2), neurocytoma (N = 2), pineoblastoma (N = 4), Rosai-Dorfman disease (N = 2), and sarcomas (N = 4). Patient sex (`Germline sex estimate`) and tumor histology (`Cancer Group`) are displayed as annotations at the bottom of each plot. Only primary tumors with mutations in the listed genes are shown. Multiple CNVs are denoted as a complex event. (B) Genome-wide plot of CNV alterations by broad histology. Each row represents one sample. Box and whisker plots of number of CNV breaks (C) or SV breaks (D) by number of chromothripsis regions.

**Figure S4:**
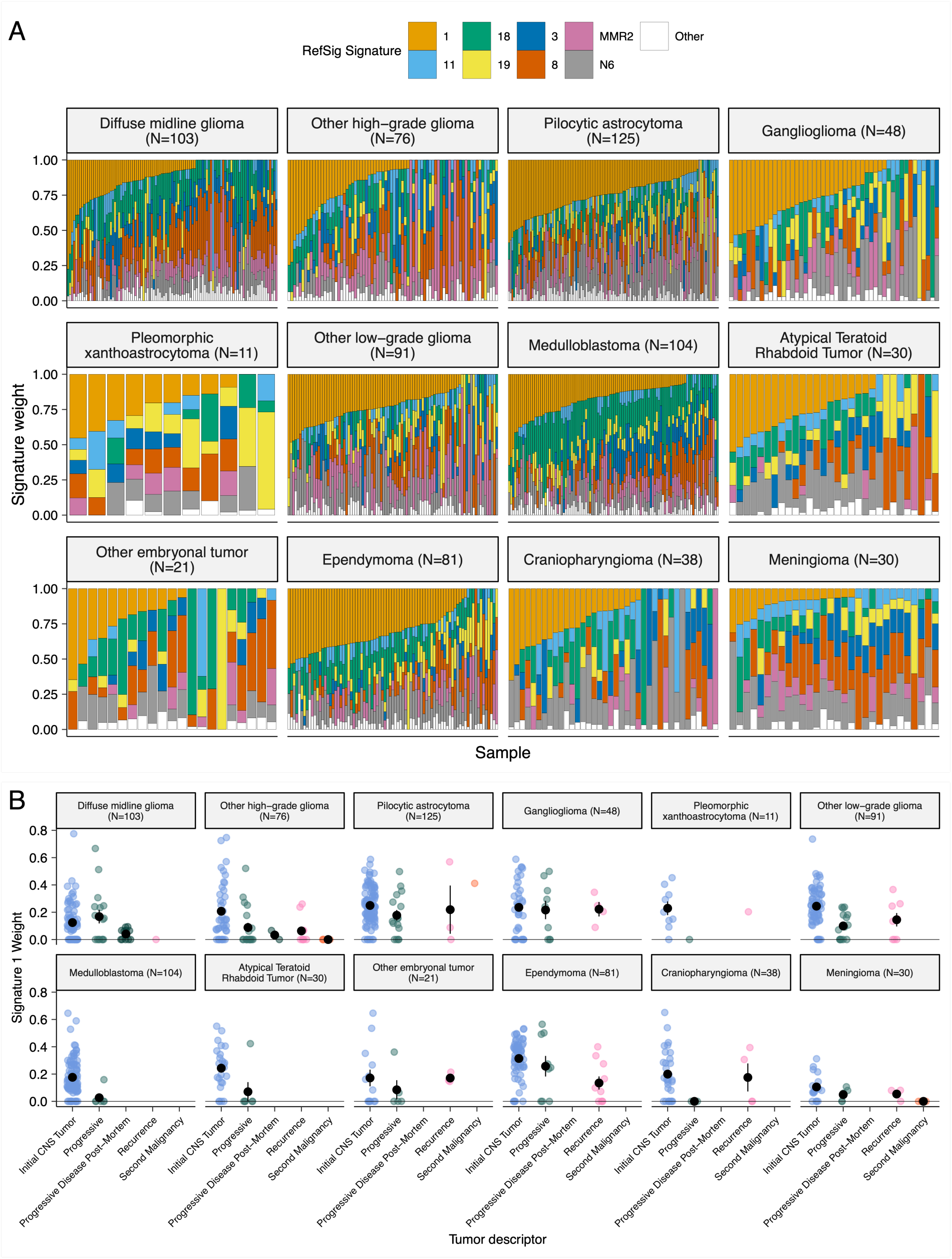
Mutational signatures in pediatric brain tumors, Related to Figure 3. (A) Sample-specific RefSig signature weights across cancer groups ordered by decreasing Signature 1 exposure. (B) Proportion of Signature 1 plotted by phase of therapy for each cancer group.

**Figure S5:**
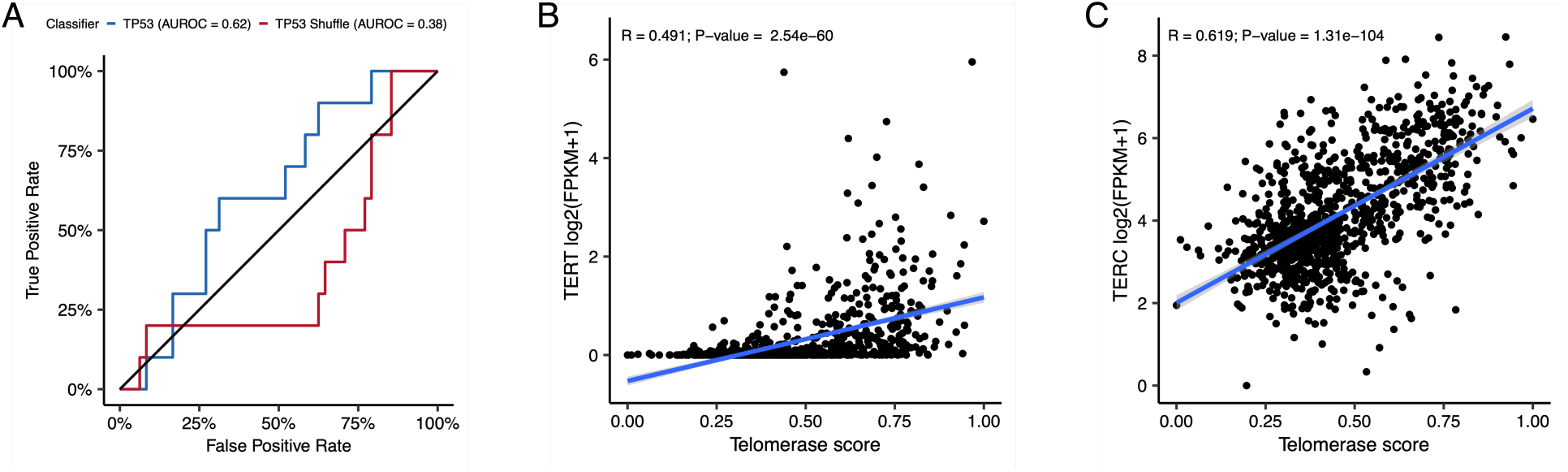
Quality control metrics for TP53 and EXTEND scores, Related to Figure 4. (A) Receiver Operating Characteristic for TP53 classifier run on FPKM of poly-A RNA-Seq samples. Correlation plots for telomerase scores (EXTEND) with RNA expression of TERT (B) and TERC (C).

**Figure S6:**
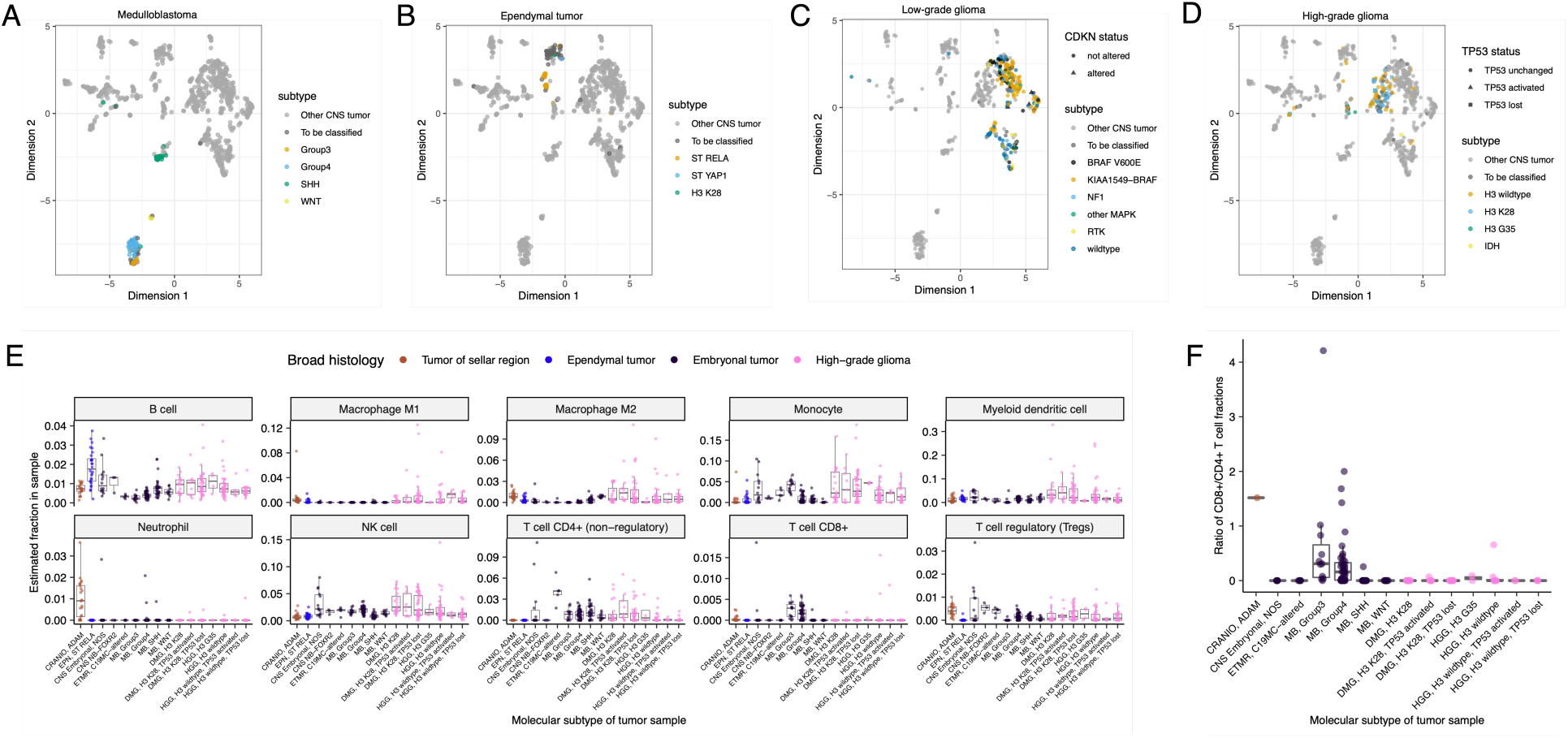
Subtype-specific clustering and immune cell fractions, Related to Figure 5. First two dimensions from UMAP of sample transcriptome data with points colored by `molecular_subtype` for medulloblastoma (A), ependymoma (B), low-grade glioma (C), and high-grade glioma (D). (E) Box plots of quanTIseq estimates of immune cell fractions in histologies with more than one molecular subtype with N >=3. (F) Box plots of the ratio of immune cell fractions of CD8+ to CD4+ T cells in histologies with more than one molecular subtype with N >=3.

**Table S1. Related to Figure 1.** Table of specimens and associated metadata, clinical data, and histological data utilized in the OpenPBTA project.

**Table S2. Related to Figures 2 and 3.** Excel file with three sheets representing tables of TMB, eight CNS mutational signatures, and chromothripsis events per sample, respectively.

**Table S3. Related to Figures 4 and 5.** Excel file with three sheets representing tables of *TP53* scores, telomerase EXTEND scores, and quanTIseq immune scores, respectively.

**Table S4. Related to Figures 4 and 5.** Excel file with six sheets representing the survival analyses performed for this manuscript. See **Star Methods** for details.

**Table S5. Related to Figure 1.** Excel file with four sheets representing of all software and their respective versions used for the OpenPBTA project, including the R packages in the OpenPBTA Docker image, Python packages i the OpenPBTA Docker image, other command line tools in the OpenPBTA Docker image, and all software used in the OpenPBTA workflows, respectively. Note that all software in the OpenPBTA Docker image was utilized within the analysis repository, but not all software was used for the final manuscript.

## Consortia

The past and present members of the Children’s Brain Tumor Network who contributed to the generation of specimens and data are Adam C. Resnick, Alexa Plisiewicz, Allison M. Morgan, Allison P. Heath, Alyssa Paul, Amanda Saratsis, Amy Smith, Ana Aguilar, Ana Guerreiro Stücklin, Anastasia Arynchyna, Andrea Franson, Angela J. Waanders, Angela N. Viaene, Anita Nirenberg, Anna Maria Buccoliero, Anna Yaffe, Anny Shai, Anthony Bet, Antoinette Price, Arlene Luther, Ashley Plant, Augustine Eze, Bailey K. Farrow, Baoli Hu, Beth Frenkel, Bo Zhang, Bobby Moulder, Bonnie Cole, Brian M. Ennis, Brian R. Rood, Brittany Lebert, Carina A. Leonard, Carl Koschmann, Caroline Caudill, Caroline Drinkwater, Cassie N. Kline, Catherine Sullivan, Chanel Keoni, Chiara Caporalini, Christine Bobick-Butcher, Christopher Mason, Chunde Li, Claire Carter, Claudia MaduroCoronado, Clayton Wiley, Cynthia Wong, David E. Kram, David Haussler, David Kram, David Pisapia, David Ziegler, Denise Morinigo, Derek Hanson, Donald W. Parsons, Elizabeth Appert, Emily Drake, Emily Golbeck, Ena Agbodza, Eric H. Raabe, Eric M. Jackson, Erin Alexander, Esteban Uceda, Eugene Hwang, Fausto Rodriquez, Gabrielle S. Stone, Gary Kohanbash, Gavriella Silverman, George Rafidi, Gerald Grant, Gerri Trooskin, Gilad Evrony, Graham Keyes, Hagop Boyajian, Holly B. Lindsay, Holly C. Beale, Ian F. Pollack, James Johnston, James Palmer, Jane Minturn, Jared Pisapia, Jason E. Cain, Jason R. Fangusaro, Javad Nazarian, Jeanette Haugh, Jeff Stevens, Jeffrey P. Greenfield, Jeffrey Rubens, Jena V. Lilly, Jennifer L. Mason, Jessica B. Foster, Jim Olson, Jo Lynne Rokita, Joanna J. Phillips, Jonathan Waller, Josh Rubin, Judy E. Palma, Justin McCroskey, Justine Rizzo, Kaitlin Lehmann, Kamnaa Arya, Karlene Hall, Katherine Pehlivan, Kenneth Seidl, Kimberly Diamond, Kristen Harnett, Kristina A. Cole, Krutika S. Gaonkar, Lamiya Tauhid, Laura Prolo, Leah Holloway, Leslie Brosig, Lina Lopez, Lionel Chow, Madhuri Kambhampati, Mahdi Sarmady, Margaret Nevins, Mari Groves, Mariarita Santi-Vicini, Marilyn M. Li, Marion Mateos, Mateusz Koptyra, Matija Snuderl, Matthew Miller, Matthew Sklar, Matthew Wood, Meghan Connors, Melissa Williams, Meredith Egan, Michael Fisher, Michael Koldobskiy, Michelle Monje, Migdalia Martinez, Miguel A. Brown, Mike Prados, Miriam Bornhorst, Mirko 1640 Scagnet, Mohamed AbdelBaki, Monique Carrero-Tagle, Nadia Dahmane, Nalin Gupta, Nathan Young, Nicholas A. Vitanza, Nicholas Tassone, Nicholas Van Kuren, Nicolas Gerber, Nithin D. Adappa, Nitin Wadhwani, Noel Coleman, Obi Obayashi, Olena M. Vaske, Olivier Elemento, Oren Becher, Philbert Oliveros, Phillip B. Storm, Pichai Raman, Prajwal Rajappa, Rintaro Hashizume, Rishi R. Lulla, Robert Keating, Robert M. Lober, Ron Firestein, Sabine Mueller, Sameer Agnihotri, Samuel G. Winebrake, Samuel Rivero-Hinojosa, Sarah Diane Black, Sarah Leary, Schuyler Stoller, Shannon Robins, Sharon Gardner, Shelly Wang, Sherri Mayans, Sherry Tutson, Shida Zhu, Sofie R. Salama, Sonia Partap, Sonika Dahiya, Sriram Venneti, Stacie Stapleton, Stephani Campion, Stephanie Stefankiewicz, Stewart Goldman, Swetha Thambireddy, Tatiana S. Patton, Teresa Hidalgo, Theo Nicolaides, Thinh Q. Nguyen, Thomas W. McLean, Tiffany Walker, Toba Niazi, Tobey MacDonald, Valeria Lopez-Gil, Valerie Baubet, Whitney Rife, Xiao-Nan Li, Ximena Cuellar, Yiran Guo, Yuankun Zhu, and Zeinab Helil.

The past and present members of the Pacific Pediatric Neuro-Oncology Consortium who contributed to the generation of specimens and data are Adam C. Resnick, Alicia Lenzen, Alyssa Reddy, Amar Gajjar, Ana Guerreiro Stucklin, Anat Epstein, Andrea Franson, Angela Waanders, Anne Bendel, Anu Banerjee, Ashley Margol, Ashley Plant, Brian Rood, Carl Koschmann, Carol Bruggers, Caroline Hastings, Cassie N. Kline, Christina Coleman Abadi, Christopher Tinkle, Corey Raffel, Dan Runco, Daniel Landi, Daphne Adele Haas-Kogan, David Ashley, David Ziegler, Derek Hanson, Dong Anh Khuong Quang, Duane Mitchell, Elias Sayour, Eric Jackson, Eric Raabe, Eugene Hwang, Fatema Malbari, Geoffrey McCowage, Girish Dhall, Gregory Friedman, Hideho Okada, Ibrahim Qaddoumi, Iris Fried, Jae Cho, Jane Minturn, Jason Blatt, Javad Nazarian, Jeffrey Rubens, Jena V. Lilly, Jennifer Elster, Jennifer L. Mason, Jessica Schulte, Jonathan Schoenfeld, Josh Rubin, Karen Gauvain, Karen Wright, Katharine Offer, Katie Metrock, Kellie Haworth, Ken Cohen, Kristina A. Cole, Lance Governale, Linda Stork, Lindsay Kilburn, Lissa Baird, Maggie Skrypek, Marcia Leonard, Margaret Shatara, Margot Lazow, Mariella Filbin, Maryam Fouladi, Matthew Miller, Megan Paul, Michael Fisher, Michael Koldobskiy, Michael Prados, Michal Yalon Oren, Mimi Bandopadhayay, Miriam Bornhorst, Mohamed AbdelBaki, Nalin Gupta, Nathan Robison, Nicholas Whipple, Nick Gottardo, Nicholas A. Vitanza, Nicolas Gerber, Patricia Robertson, Payal Jain, Peter Sun, Priya Chan, Richard S Lemons, Robert Wechsler-Reya, Roger Packer, Russ Geyer, Ryan Velasco, Sabine Mueller, Sahaja Acharya, Sam Cheshier, Sarah Leary, Scott Coven, Sebastian M. Waszak, Sharon Gardner, Sri Gururangan, Stewart Goldman, Susan Chi, Tab Cooney, Tatiana S. Patton, Theodore Nicolaides, and Tom Belle Davidson.

